# Clonally resolved single-cell multi-omics identifies routes of cellular differentiation in acute myeloid leukemia

**DOI:** 10.1101/2022.08.29.505648

**Authors:** Sergi Beneyto-Calabuig, Anne Kathrin Ludwig, Jonas-Alexander Kniffka, Chelsea Szu-Tu, Christian Rohde, Magdalena Antes, Alexander Waclawiczek, Sarah Gräßle, Philip Pervan, Maike Janssen, Jonathan J. M. Landry, Vladimir Benes, Anna Jauch, Michaela Brough, Marcus Bauer, Birgit Besenbeck, Julia Felden, Sebastian Bäumer, Michael Hundemer, Tim Sauer, Caroline Pabst, Claudia Wickenhauser, Linus Angenendt, Christoph Schliemann, Andreas Trumpp, Simon Haas, Michael Scherer, Simon Raffel, Carsten Müller-Tidow, Lars Velten

**Affiliations:** Centre for Genomic Regulation (CRG), The Barcelona Institute of Science and Technology, Dr. Aiguader 88, Barcelona 08003, Spain; Department of Medicine, Hematology, Oncology and Rheumatology, University Hospital Heidelberg, Heidelberg, Germany; Universitat Pompeu Fabra (UPF), Barcelona, Spain; Molecular Medicine Partnership Unit, European Molecular Biology Laboratory (EMBL), University of Heidelberg, Heidelberg, Germany; Heidelberg Institute for Stem Cell Technology and Experimental Medicine (HI-STEM gGmbH), Heidelberg, Germany; Division of Stem Cells and Cancer, Deutsches Krebsforschungszentrum (DKFZ) and DKFZ-ZMBH Alliance, Heidelberg, Germany; Berlin Institute of Health (BIH) at Charité – Universitätsmedizin Berlin, 10117 Berlin, Germany; Charité-Universitätsmedizin, 10117 Berlin, Germany; Berlin Institute for Medical Systems Biology, Max Delbrück Center for Molecular Medicine in the Helmholtz Association, 10115 Berlin, Germany; Genomics Core Facility, European Molecular Biology Laboratory (EMBL), Heidelberg, Germany; Institute of Human Genetics, University of Heidelberg, Heidelberg, Germany; Institute of Pathology, University Hospital Halle (Saale), Martin-Luther-University Halle-Wittenberg, 06112 Halle, Germany; Department of Medicine A, Hematology and Oncology, University Hospital Muenster, Germany; Department of Biosystems Science and Engineering, ETH Zurich, Basel, Switzerland

## Abstract

Inter-patient variability and the similarity of healthy and leukemic stem cells have impeded the characterization of leukemic stem cells (LSCs) in acute myeloid leukemia (AML), and their differentiation landscape. Here, we introduce CloneTracer, a novel method that adds clonal resolution to single-cell RNA-seq datasets. Applied to samples from 19 AML patients, CloneTracer revealed routes of leukemic differentiation. While residual healthy cells dominated the dormant stem cell compartment, active leukemic stem cells resembled their healthy counterpart and retained erythroid capacity. By contrast, downstream myeloid progenitors were highly aberrant and constituted the disease-defining compartment: Their gene expression and differentiation state determined both chemotherapy response and the leukemia’s ability to differentiate to transcriptomically normal monocytes. Finally, we demonstrated the potential of CloneTracer to identify surface markers mis-regulated specifically in leukemic cells by intra-patient comparisons. Taken together, CloneTracer revealed a differentiation landscape that mimics its healthy counterpart and determines biology and therapy response in AML.

## Introduction

Our understanding of blood formation has fundamentally changed in the last decade. In particular, single-cell RNA-seq based studies have demonstrated that hematopoietic stem cells acquire priming early, at phenotypically immature stages^1^. The oligopotent progenitor types that were previously thought to drive hematopoiesis, such as common myeloid progenitors (CMPs), consist of mixtures of fully lineage committed cells^2^. Rather than passing through a CMP stage, lineage differentiation occurs along two major branches, a lymphomyeloid and an erythromyeloid branch^3, 4^. These results are supported by various functional assays^4–7^. By contrast, the aberrations that characterize the differentiation landscape in myeloid malignancies remain unknown. In particular, many myeloid malignancies were thought to affect, or originate from, CMPs, which do not represent a defined cell type. Knowledge of differentiation tracks of malignant cells might enable novel diagnostic and therapeutic approaches.

Since the healthy and the diseased hematopoietic system co-exist in myeloid malignancies, investigating malignant differentiation landscapes requires clonally resolved single-cell RNA-seq methods. Recent studies have profiled CALR-mutant^8^ and JAK2-mutant^9^ myeloproliferative neoplasm, as well as DNMT3A-mutant clonal hematopoiesis^10, 11^ and revealed the expansion of particular differentiation states at the expense of others, and a “skewing” of the hematopoietic differentiation landscape^12^. However, the shape of the cellular differentiation landscape in full-blown leukemia such as acute myeloid leukemia (AML) remains unknown: Are healthy routes of lineage differentiation co-opted in this disease, or are novel, aberrant cellular identities created? And, more specifically, are leukemic stem cells (LSCs) a consistent cell type resembling healthy HSCs, or are they heterogeneous groups of leukemic cells that possess certain stemness properties? Answering these questions is of key importance to prioritize cellular targets of therapies, identify novel prognostic factors, and identify pathways altered in cancer, but not in healthy cells of a similar differentiation state.

To answer these questions, we have developed a new strategy to add clonal resolution to high-throughput (droplet-based) single-cell RNA-seq data that robustly works across many of the heterogeneous AML genotypes. Existing approaches use single nucleotide variant (SNVs) or mitochondrial SNVs (mtSNVs) as qualitative markers to identify healthy and malignant cells from scRNAseq data^8–10, 13, 14^. However, these measurements are noisy and biased by cellular identity impeding quantitative analysis. Copy number variants (CNVs) can be inferred from scRNAseq data^15–18^, but are not always present in AML and other leukemias. Our new computational method, CloneTracer, integrates information from SNVs, mtSNVs and infers CNVs (when present) through a statistical model that appropriately accounts for the noise properties of these measurements. CloneTracer thereby identifies the clonal hierarchy for each sample and probabilistically assigns single cells to clones. CloneTracer overcomes limitations of previous computational methods that either were designed for single-cell DNA-seq data^19–21^, or require prior knowledge about the clonal hierarchy^22^.

We applied CloneTracer to bone marrow samples from 19 patients with AML; four of these patients included follow-up samples. We showed that CloneTracer could unanimously identify most healthy and leukemic cells in 13 of these patients. By integrating data across all patients, we identified a population of HSCs expressing a dormancy gene signature that was dominated by residual healthy cells, as well as leukemic cells resembling active HSCs (aLSCs) that often retained erythroid capacity. Downstream of aLSCs, differentiation-blocked, aberrant myeloid progenitors determined chemotherapy response and fed into qualitatively normal myeloid differentiation at rates that could be determined from their transcriptome. Together, our data established a healthy-like differentiation landscape that determines biology and therapy response in leukemia.

## Results

### A method for optimizing the single-cell coverage of nuclear and mitochondrial SNVs in droplet-based single-cell RNA-seq

Conventional droplet based single-cell RNA-seq protocols exhibit low coverage for nuclear and mitochondrial mutations. Hence, we first sought to expand the mutational information available from 10x genomics single cell RNA-seq libraries. Following cDNA amplification by the default CITE-seq^23^ protocol, the cDNA pool was split and sequencing libraries specifically covering RNA expression, surface antigen expression, nuclear SNVs and mitochondrial genomes were generated (Figure 1a, methods). To cover mitochondrial genomes at full-length, we used PCR primers located near the 5’ end of mitochondrial genes, and subsequently performed an optimized tagmentation protocol (Figure S1a,b). Full length coverage was substantially improved compared to default 10x, and similar to a recent report that optimized mitochondrial coverage using a purely PCR-based strategy^14^ (Figure 1b). Overall, 55-85% of these libraries mapped to the mitochondrial genome, allowing for a cost-effective deep sequencing of mitochondrial genomes. A cell line mixing experiment illustrated that false positive observations of mitochondrial genetic variants occurred at negligible background rates (Figure S2).

**Figure 1.**
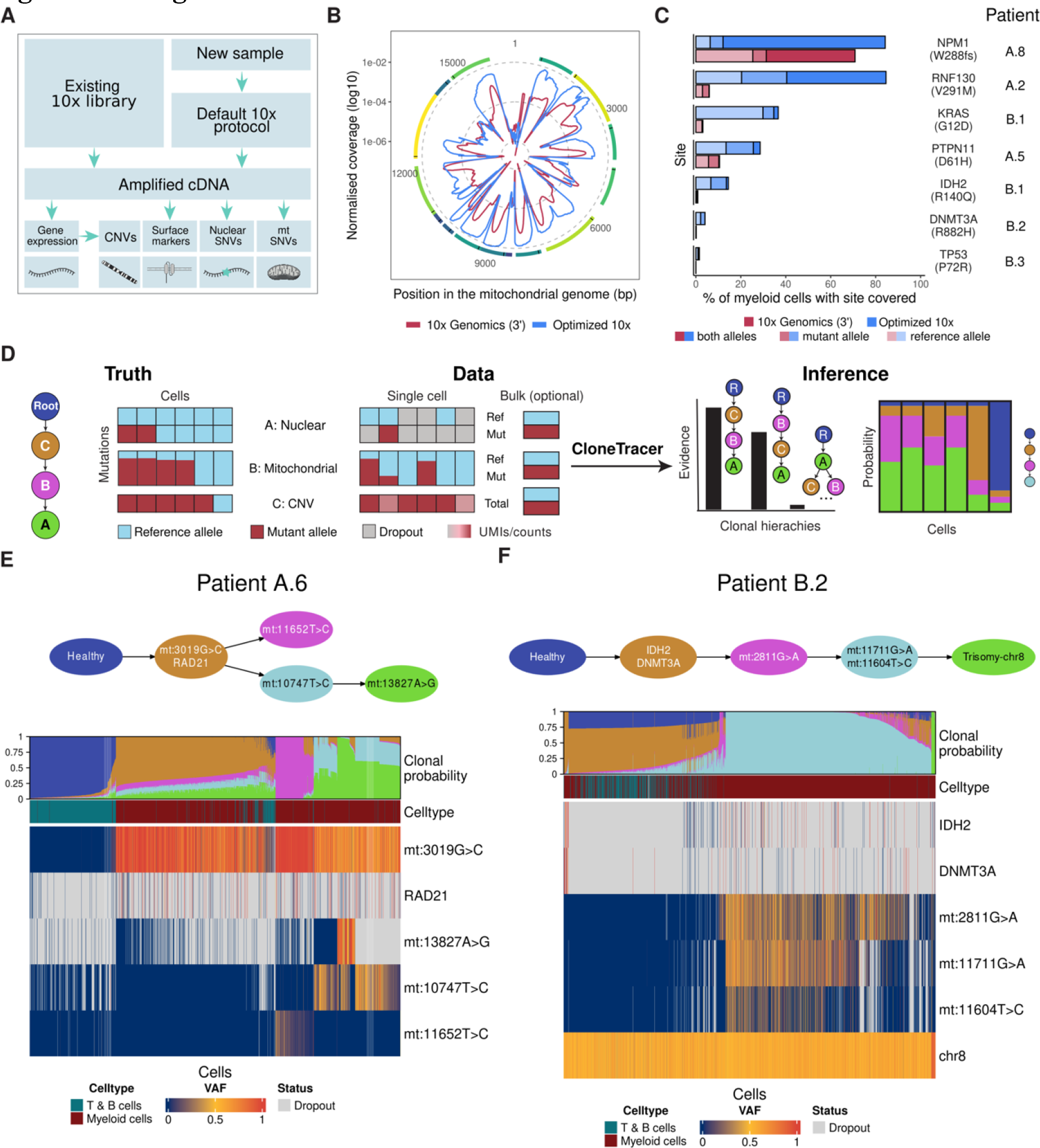
CloneTracer and “Optimized 10x” enable clonal tracking in droplet-based single-cell RNA-seq. **a.** Scheme of “Optimized 10x”. **b.** Normalized coverage across the mitochondrial genome obtained by baseline and Optimized 10x. **c.** Coverage of nuclear mutations from various AML patients (see figure 2) profiled with CloneTracer or baseline 10x genomics. Only immature and early myeloid cells are included. **d.** Illustration of the statistical challenge addressed by CloneTracer. In this example, a patient displays a sub-clonal nuclear mutation, a mitochondrial mutation and a CNV. For the nuclear mutation, two cells carry a mutated allele while four cells carry two healthy alleles. For the mitochondrial mutation, four cells carry the mutation on a fraction of their mitochondrial genomes while two cells do not display the mutation. For the CNV, five cells display the copy number variant. This ground truth is observed through a noisy single cell RNA-seq process, which introduces false negatives and false positives, and in some cases, data is missing. Additionally, bulk allele frequencies or clonal fractions are observed. The goal of the model is to infer the hierarchy among the clones, and probabilistically assign each cell to a clone while accounting for the measurement errors. **e.** Top row: Inferred clonal hierarchy for patient A.6. Middle row: Stacked bar chart illustrating each cell’s probability to derive from the different clones shown in top panel. Bottom rows: Heatmap depicting the variant allele frequency of all clonal markers in all cells. **f.** Like panel e, except that data from patient B.2 is shown.

For genotyping of nuclear SNVs, we utilized a modified version of TAP-seq^24^ which includes 3 nested PCR steps with mutation-specific primers (Figure S1c-e, and https://github.com/veltenlab/CloneTracer/tree/master/primer_design for a primer design software that also assists with identifying mutations suitable for amplification). Thereby, coverage on disease relevant mutations was substantially improved, compared to baseline 10x genomics 3’ (Figure 1c), and similar to results reported by others with related approaches^8, 9^. Importantly, mitochondrial and nuclear SNV targeted sequencing libraries could readily be constructed from existing (e.g., already sequenced) full length cDNA libraries from 10x genomics (step 2.3A), making this method (“Optimized 10x”) applicable to characterize existing samples with more depth. Overall, 10ng of cDNA were sufficient to prepare both mitochondrial and nuclear SNV libraries.

We compared the performance of “Optimized 10x” to related methods, a plate-based RNA-seq protocol (MutaSeq, a modified Smart-Seq2 protocol^25^) and a droplet-based ATAC-seq method focused on tracking mitochondrial mutations (sc-mito-ATAC-seq^26^) (Figure S3). In sum, “Optimized 10x” maximizes the mutational information available from default 10x genomics libraries (Figure S3A,B), and, unlike low-throughput high-confidence plate-based methods^25, 27^, it has the throughput required for ambitious single cell RNA-seq oncology projects (Figure S3C).

### CloneTracer, a statistical model to infer clonal hierarchies and clonal assignments for single-cell RNA-seq based SNV, mitochondrial SNV and CNV data

Despite the improved coverage of leukemic point mutations and mitochondrial SNVs in “Optimized 10x”, the data from RNA-seq based protocols is always noisy, as illustrated by frequent allelic dropout even of highly expressed genes such as *NPM1* (see Figure 1c, S3). For a confident interpretation of the data and quantitative analyses, statistical methods are needed that identify the most likely hierarchy among the mutations and thereby, for example, clarify if a mitochondrial mutation or CNV marks all cancer cells or merely a subclone. Further, dropout and false positive rates need to be systematically accounted for when assigning cells to clones.

We therefore developed CloneTracer, a Bayesian model that identifies the hierarchical relationship between mutations and exploits this hierarchy when assigning cells to clones. Our model considers any prior information, such as allele frequencies from bulk exome sequencing, and, most importantly, it accounts for the technical noise associated with single-cell mutation measurements of CNVs, SNVs and mtSNVs (Figure 1d and see supplementary methods for a detailed description of the model). Our model achieved two main tasks: First, it compared possible clonal hierarchies and computed the most likely configuration of mutations across the cells given the data. Second, for the mutational hierarchy with the highest evidence, it computed the posterior probability of each cell to belong to any particular clone.

To demonstrate the performance of CloneTracer, we first highlight two patients from our 19-patient cohort (see below for a detailed description of the cohort). In the case of patient A.6, a nuclear SNV with a high allele frequency in bulk exome-seq was located in *RAD21*, which was covered in 13.3% of single cells. However, its mutant allele was only observed in cells carrying also at least one mitochondrial mutation. Hence, cells could be identified as healthy or leukemic, with a confidence that mostly depended on the coverage of the mitochondrial SNVs in each cell (Figure 1e, and see Supplementary Table 3 for exome data).

In the case of patient B.2, nuclear SNVs with a high VAF were identified in *IDH2* and *DNMT3A*. While these mutations mostly co-occurred with mitochondrial mutations, they were also occasionally observed on their own, indicating that here, the mitochondrial mutation occurred downstream of the first leukemogenic hit (Figure 1f). Hence, in this patient, most leukemic cells were identified with a high confidence, but a confident assignment of healthy vs. *DNMT3A/IDH2* mutant early leukemic cells was not possible. Together, these examples illustrate the importance of using statistical models when interpreting noisy single cell genotyping data.

### Application of CloneTracer to primary AML patient specimens

In total, we applied CloneTracer to 19 AML patients from two study cohorts (Figure 2a,b, Supplementary Table 3). Cohort A consisted of diagnostic bone marrow samples that were characterized by whole exome sequencing and subjected to a CD34 enrichment prior to single cell CITE-seq; a median of 2,232 single cells per patient passed stringent quality control filters. Cohort B included sequential samples from four individual patients at times of diagnosis, after therapy and (in one case) relapse samples. Genotypes were analyzed by panel sequencing and cells were not subjected to enrichment prior to single cell CITE-seq. A median of 12,034 single cells per patient passed stringent quality filters. In both cohorts, cells were stained with CITE-seq surface antibodies prior to sequencing (37 antibodies in cohort A, 79 antibodies in cohort B, see Supplementary Table 1). Overall, we analyzed 88,602 single cells from 25 individual specimens.

**Figure 2.**
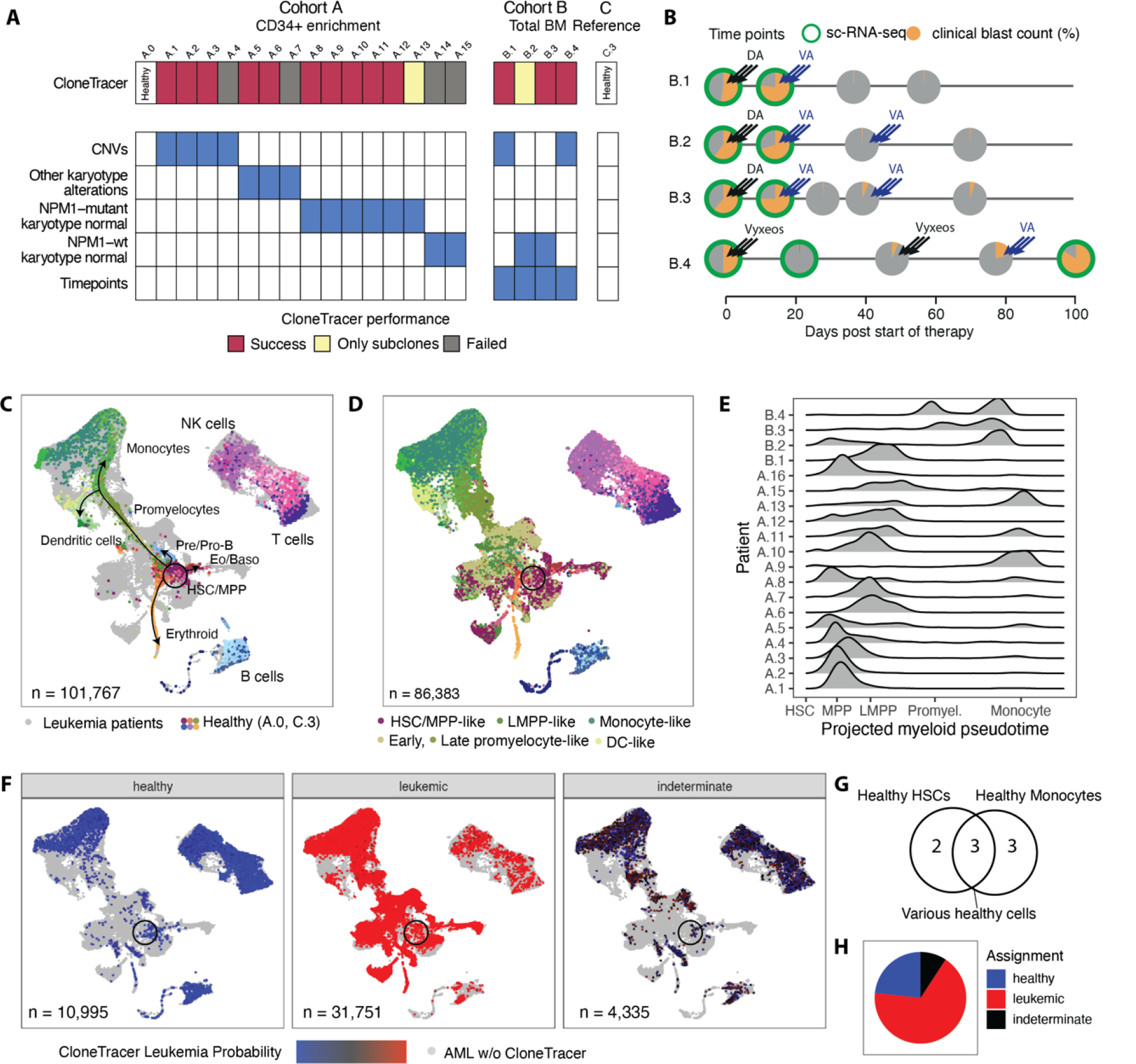
CloneTracer applied to two AML patient cohorts. **a.** Overview of genotypes and CloneTracer performance. **b.** Overview of longitudinal sampling in cohort B. Pie charts indicate clinical blast counts at the respective BM aspiration time points. DA/VA indicate treatment with Daunorubicin/Ara-C and Venetoclax/Azacitidine, respectively. Of note, clinical blast count of patient B.1 and B.3 conflicts with scRNA-seq estimate, see Figure S4b. **c.** uMAP depicting integrated data from both cohorts. Colors (see also Figure S4a) highlight cell states of cells from two healthy individuals (A.0 and C.3^4^). Cells from leukemia patients are shown in grey. **d.** Cells from leukemia patients were projected to the healthy reference^4^ to identify the most similar healthy cell type and corresponding pseudotime of myeloid differentiation for every cell, see methods for detail. Panel d highlights projected cell types on an uMAP, see figure S4a for a complete color legend. **e.** Density chart of cells over projected pseudotime, stratified by patient. Density was corrected to account for FACS enrichment in cohort A: Fraction of total bone marrow is shown. For cohort B, only data from day 0 is included. **f.** uMAP highlighting CloneTracer leukemia probabilities for 13/19 AML patients. Cells from the remaining individuals are shown in grey. For cohort B, only data from day 0 is included. **g.** Venn diagram highlighting the patients with contribution of healthy monocytes and healthy HSCs. **h.** Pie chart summarizing the assignment of cells as healthy or leukemic across the 13 patients.

Clonal hierarchies were identified as described for the examples of patient A.6 and B.2. A detailed description of this analysis is available in the supplementary note; in summary, CloneTracer was able to unanimously identify healthy and leukemic cells in 13/19 patients, including A.6 (Figure 2a). Of note, it is possible that some cells assigned as healthy might carry pre-leukemic mutations, for example in *DNMT3A* or *TET2*. In two additional patients, including B.2, many cancer cells were unanimously identified as such but a confident assignment of healthy vs. early leukemic cells was not possible. In the remaining patients, most cells cannot be assigned to cancer or healthy with high confidence, due to a lack of highly covered clonal markers. Unless explicitly stated otherwise, all subsequent analyses that take clonal identities into account pertain to the 13 patients with high confidence healthy/leukemic assignments.

To identify differentiation landscapes, we integrated the data from all 19 patients with data from a healthy individual processed as part of cohort A (A.0) as well as a second reference healthy individual for which detailed cell state annotation was available (C.3, from ref. ^4^). This integration identified clusters of lymphoid cells, monocytes, dendritic cells (Figure 2c), as well as a highly heterogeneous group of immature myeloid leukemic progenitors. Each cell was projected onto the healthy reference^4^ in order to identify the most similar healthy cell state (Figure 2d, S4a). We also assessed a corresponding pseudotime value of myeloid differentiation (Figure 2e). Highlighting the healthy vs. leukemia assignments of CloneTracer on this map demonstrated that lymphoid cells were mostly assigned as healthy whereas myeloid cells were mostly assigned as leukemic (Figure 2f, S4b, and see Figure 3a for quantification). Of note, healthy monocytes and HSC/MPP-like cells occurred in several patients (Figure 2g, S4b). We then assigned all cells as leukemic if their posterior probability for being leukemic was >0.8, healthy if the posterior probability was <0.2, and excluded them from clonal analyses otherwise. Overall, 91% of cells from the 13 patients could thereby be assigned as healthy or leukemic (Figure 2h).

**Figure 3.**
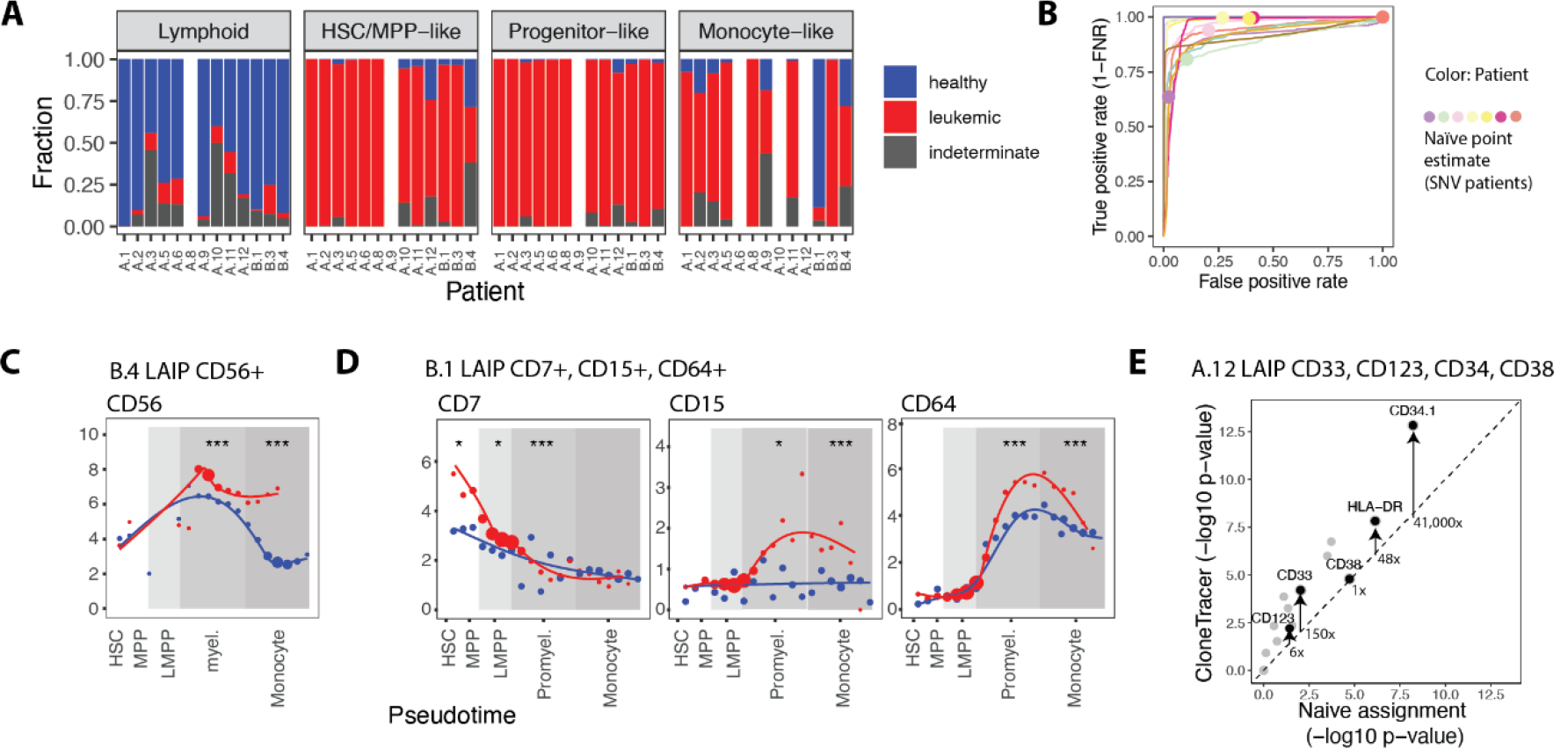
Validation of CloneTracer clonal assignments. **a.** Barchart depicting the fraction of cells assigned as healthy (blue), leukemic (red) or indeterminate (grey), stratified by projected cell type. Cell types not covered with at least 50 cells in a patient are omitted. **b.** ROC curves computed from CloneTracer leukemia probabilities, using the ground truth that lymphoid cells are healthy and all myeloid cells are leukemic. Dots depict statistically naïve point estimates for the patients without CNVs. **c.** Smoothened expression of LAIP markers over projected pseudotime of patient B.4, stratified by clone. Points indicate mean expression within 20 equally sized bins along pseudotime, point size indicates number of cells per bin. Asterisks indicate significance of differential expression within the shaded area of pseudotime. ***: FDR < 0.001, **: FDR < 0.01, *: FDR < 0.1. p values are from a wilcoxon test of library-size normalized ADT counts. **d.** As in c, but for patient B.1. **e.** Scatter plot depicting p-values from statistical tests comparing surface marker expression between healthy and leukemic immature cells. x axis shows estimates obtained using a statistically naïve assignment, y axis shows estimates obtained using CloneTracer assignments.

### Validation of CloneTracer and comparison to naïve clonal assignments

We validated the assignments using established parameters for AML diagnosis.

A regular assumption of AML diagnosis is that most myeloid cells should be leukemic, whereas lymphoid cells should be healthy. Under this assumption, the median area under the receiver-operating characteristics curve (AUROC) of CloneTracer was 0.96 (range 0.88-1). The discretized CloneTracer assignments had a median false positive rate (FPR) across patients of 9% (range 0.1%-20%), and a median false negative rate (FNR) of 1% (range 0%-20%, Figure 3a,b). For patients with SNVs as clonal markers, statistically naïve assignments^8, 28^ that classify a cell as leukemic if at least one mutant allele is observed, and otherwise as healthy if at least one healthy allele is observed, in most cases sub-optimally balanced between FPR and FNR (Figure 3b).

Notably, not all myeloid cells are really leukemic. We therefore considered “Leukemia Associated Immunophenotypes” (LAIPs) to distinguish leukemic vs. healthy myeloid cells. LAIPs are cell state specific, aberrantly expressed markers identified during routine clinical flow cytometric analysis of AML diagnostic samples^29, 30^. Three patients carried a significant number of residual healthy cells along the full myeloid differentiation spectrum, as well as a clinically described LAIP. As expected, leukemic, but not residual healthy cells, expressed the LAIP in a differentiation-state dependent manner (Figure 3c,d). For the patient with described LAIP and SNVs as clonal marker, the statistical power in identifying LAIP markers was increased by using CloneTracer, compared to the statistically naïve assignment (Figure 3e).

Together, these analyses demonstrate that CloneTracer correctly assigns cells as healthy and leukemia and outperforms a statistically naïve assignment.

### Differentiation hierarchies in leukemia

We next utilized the high confident healthy/leukemia assignments to investigate differentiation routes in AML. We aimed to identify leukemic and healthy stem cells and to visualize their downstream paths of differentiation. Of note, one cluster of cells, as defined from gene expression (“C6”, Figure 4a, S5a), contained HSCs from the healthy reference individuals, as well as healthy and leukemic cells from the AML patients (see also Figure 2c,f). Transcriptomes suggested an HSC and early MPP identity (see also Figure 2d). We also computed the average transcriptomic similarity to the five most similar healthy cells, conceptually an aberration score, and observed that cells from C6 were relatively similar to HSC/MPPs, whereas other immature progenitors were more aberrant (Figure 4b).

**Figure 4.**
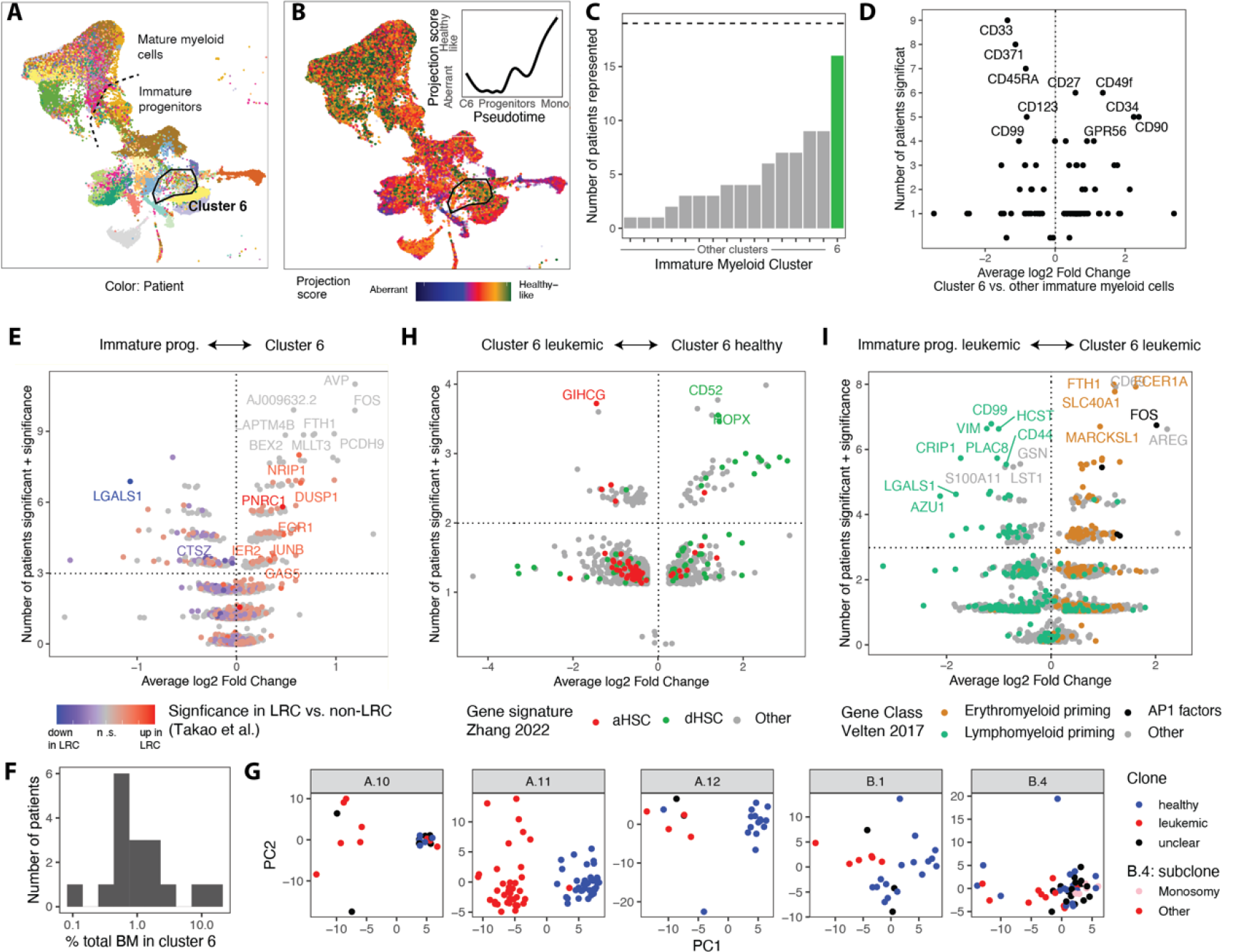
Identification and characterization of LSCs. **a.** uMAP of n = 56,748 myeloid cells from the 19 leukemia patients, highlighting the patient identity. **b.** uMAP highlighting the projection score, i.e. similarity to the 5 most similar healthy reference cells, see also methods. Inset depicts a smoothened average of the projection score as a function of pseudotime. **c.** Bar chart summarizing the number of patients represented in each cluster with at least 5 cells. n=19 patients were analyzed. **d.** Volcano plot highlighting the number of patients where a given surface marker was significantly (p < 0.05) differentially expressed between cells from C6 and other immature myeloid cells from the same patient, vs. the average log2 fold change across patients. n=16 patients with C6 represented and n=2 healthy individuals were analyzed. **e.** Volcano plot as in d, but for RNA expression. On the y axis, the number of patients where the gene is differentially expressed was added to the scaled sum of log p-values from the individual comparisons. The log sum of p values was scaled to a maximum of 1. Genes are colored using information on their expression in label retaining (LRC) vs. non label retaining AML cells^31^. Darker colors indicate that the gene appeared as significant in LRC-non-LRC comparisons from more patients. **f.** Histogram depicting the size of C6 as a fraction of total bone marrow. n=16 patients with C6 are included. **g.** RNA expression data of cells from C6 were separately analyzed for each patient to normalize the data, identify highly variable genes and compute principal components (PCs). PC score plots are shown separately by patient, highlighting CloneTracer assignments. n=5 patients with both healthy and leukemic cells represented in C6 are shown, see also Figure S5c. **h.** Volcano plot as in e, comparing healthy and leukemic cells from C6. n=5 patients were analyzed, as in panel g. Genes are colored by human HSC gene signatures^32^. **i.** Volcano plot as in panel e, comparing leukemic cells from C6 to other leukemic immature myeloid cells from the same patient. n=13 patients with confident CloneTracer leukemia assignments were analyzed. Genes are colored by priming gene signatures^1^.

Interestingly, C6 contained cells from 16 out of 19 AML patients and both healthy individuals, whereas most other progenitor clusters were dominated by cells from only one patient each (Figure 4a,c). Compared to other progenitor cells, cells from cluster 6 tended to have a stem cell surface marker profile (Figure 4d: high level expression of CD34, CD90, lower expression of CD38, in leukemia patients, additionally increased expression of GPR56). Interestingly, compared to other progenitors, C6 expressed genes identified as upregulated in label-retaining AML cells (LRCs) during xenotransplant assays, compared to the non-label-retaining fraction (Takao & Kentsis, submitted and ref. ^31^), further underlining the idea that C6 contains a leukemia stem cell population (Figure 4e, Supplementary Table 2). LRCs were characterized as the population responsible for causing disease in xenotransplants, and for drug resistance (Takao & Kentsis, submitted). Across the different AML patients, cluster 6 contained a median of 1% (range 0.1%-17%) of total bone marrow (Figure 4f) which is somewhat higher than the LSC number estimated from xenotransplants.

Leukemic cells outside of C6 mostly resembled downstream myeloid progenitor states (see also Figure 2d). Cells clustered in patient-specific manner and displayed gene expression profiles rather dissimilar to healthy myeloid progenitors, highlighting the large inter-patient heterogeneity of AML (Figure 4a,b). At the level of monocytes and dendritic cells, the transcriptomic similarity between leukemic cells from various patients and healthy cells increased (Figure 4a,b). We therefore conceptually structure leukemic differentiation in three stages, putative healthy-like stem cells (“C6”), highly heterogeneous and aberrant progenitors, and mature, healthy-like monocytes/dendritic cells.

### The dormant stem cell compartment is predominantly healthy

To characterize C6, we first focused on the five patients for whom both healthy and leukemic cells occurred within this cluster. With very few exceptions, healthy and leukemic cells in C6 separated by principal component analysis based on gene expression (Figure 4g). Healthy cells expressed genes characterized as “dormant HSC” genes in 1) a recent single-cell RNA-seq study of highly purified human HSCs^32^, 2) label retention assays in mice^33^, and 3) “low output HSC” genes identified using clonal tracking^34^ (Figure 4h, S5b, Supplementary Table 2). The dormant HSC gene expression signature robustly separated healthy from the large majority of leukemic C6 cells across patients (Figure S5c); cells expressing the dormant signature were consistently CD34+CD38- (Figure S5d). These results suggest that in AML, the dormant stem cell compartment, where present or observed, contains healthy stem cells (dHSCs).

### Leukemic stem cells with erythro-myeloid priming and erythroid capacity are common in AML

Transcriptomes of the leukemic fraction in C6 exhibited increased variability between patients, compared to the predominantly healthy, putatively dormant fraction. In some patients, leukemic cells from C6 expressed “active HSC” genes^32, 33^ or “high output HSC” genes^34^ relative to healthy cells from C6 (Figure 4h, S5b, Supplementary Table 2). To identify if these cells are truly distinct from other leukemic progenitors we performed differential expression testing, contrasting leukemic C6 cells with other leukemic myeloid progenitor cells from the same patient. In most patients, leukemic C6 cells highly expressed genes associated with early erythromyeloid (erythroid, megakaryocytic and eosinophilic/basophilic) priming^1^ as well as AP-1 transcription factors such as *FOS* (Figure 4i, S5e, Supplementary Table 2). By contrast, leukemic cells from the other myeloid progenitor clusters expressed genes associated with lymphomyeloid priming^1^ and lacked the expression of consistent erythromyeloid priming signatures. Of note, genes overexpressed by leukemic cells from C6, compared to other progenitors, were also enriched (p<10^-12^) for genes associated with label retaining AML cells in xenotransplants (Takao & Kentsis, submitted) (Figure S5f, Supplementary Table 2).

Of note, in 10/13 patients with confident CloneTracer assignments, we observed erythroid progenitors carrying leukemic mutations. In many patients these cells carried NPM1 mutations and/or CNVs, and were hence not pre-leukemic, but actually derived from the leukemic clone. The abundance of mutation carrying erythroid progenitor cells correlated with the abundance of C6 (Figure 5a, see also S4b). These results indicate that most leukemias, and specifically C6, can differentiate into the erythroid lineage.

**Figure 5.**
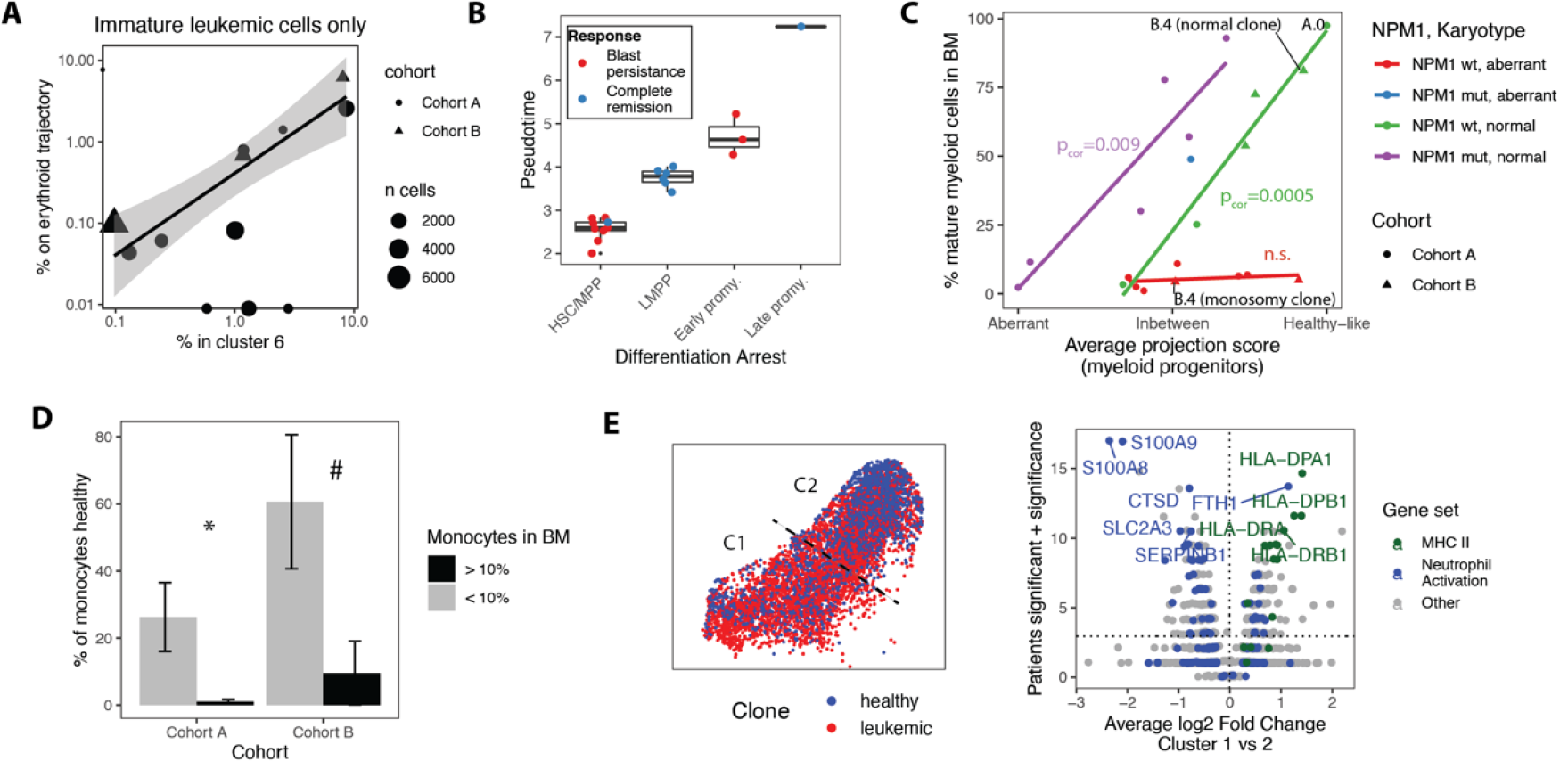
Differentiation pathways downstream of LSCs. **a.** Scatter plot depicting the fraction of immature leukemic cells falling in cluster 6 vs. the fraction of immature leukemic cells mapped on the erythroid trajectory (cluster 29). n=13 patients. **b.** Boxplot contrasting the therapy response of the patient with the average pseudotime of the immature leukemic cells. **c.** Scatter plot depicting the fraction of mature myeloid cells in diagnostic bone marrow samples across n=19 leukemic and one healthy individual (A.0) as a function of the average projection score of the immature myeloid progenitors. Genotype is color coded. For Patient B.4, the fraction of mature myeloid cells was computed separately for the two subclones. **d.** Bar chart depicting the fraction of healthy monocytes as a function of the total monocyte abundance. Only patients with confident CloneTracer assignments were analyzed: n=10 patients for cohort A and n=7 sampling timepoints for cohort B. **e.** Left panel: CloneTracer assignments on the uMAP of cluster 1&2 (monocytes), n=5,818 cells. Right panel: Volcano plot as in figure 4e, comparing monocyte cluster C1 to monocyte cluster C2 from the same patient. n=19 patients and n=1 healthy individuals were analyzed.

Together these results justify to designate leukemic cells from C6 as active Leukemic Stem Cells (aLSCs). The abundance of aLSCs did not correlate with the stage and degree of the downstream differentiation block of leukemic myeloid progenitors. Accordingly, aLSCs are a compartment distinct from the lympho-myeloid progenitors that make up most of the disease. In sum, aLSCs are a rare population of stem-like cells that exists in most AML patients and that often retains erythroid capacity.

### The state and extent of the differentiation block determines the phenotypic manifestation of AML

Leukemic progenitors downstream of LSCs can be projected onto healthy progenitor states, ranging from stem cells to promyelocytes^4, 28^ (see also Figure 2c-e). The average projected pseudotime predicted therapy response (Figure 5b): The patients with the most immature leukemic progenitors had blast persistence or died during induction therapy whereas patients with LMPP-like leukemic progenitors went into complete remission following first-line chemotherapy (p=0.001, Fisher test). The few cases with an arrest at promyelocyte stage displayed a trend towards blast persistence in the case of an earlier differentiation arrest. The presence or size of the C6 stem cell population was not correlated with first-line chemotherapy response.

These results suggest that the stage of the differentiation block is an important property of the AML, and possibly subject to evolutionary selection. We therefore investigated three patients where CloneTracer had confidently identified co-existing sub-clones marked by relevant driver mutations. In patient B.1, we observed three clones: A trisomy 8/IDH2 mutant clone, as well as a KRAS and a NRAS mutant subclone. While all clones existed both in a GPR56-high, more immature state, and a GPR56-low, late promyelocyte-like state, the KRAS clone was predominantly observed in the more immature state (Figure S6a-c). In patient B.4, we observed two clones, one of which was marked by monosomy 7, and the other one by a mitochondrial mutation. According to bulk exome data, both clones carried a TET2 mutation. While the mitochondrial clone abundantly generated monocytes, the monosomy 7 clone existed in a promyelocyte-like state and relapsed after 105 days while maintaining its cell state (Figure S6d). In patient A.2, a monosomy 7 clone co-existed with a -7/+8 subclone in a patient-specific aberrant cell state. Within this state, the subclone shifted to a more immature identity and altered its cell surface phenotype from GMP-like to MEP-like (Figure S6g). Across the full cohort, the presence of subclones, as identified from bulk exome sequencing, was associated with a larger variability in terms of differentiation status, i.e. projected pseudotime (Figure S6h). Taken together, in the three cases investigated, subclones were shifted towards more immature differentiation blocks, compared to the parental clones, possibly because evolutionary pressures favor differentiation blocks at more immature states.

We next investigated the ability of the leukemic progenitors to give rise to mature monocytes. As introduced before, leukemic SCs, and monocytes/dendritic cells were rather healthy-like, whereas leukemic progenitors displayed variable degrees of aberration across patients (see figure 4b). We found that this aberration score correlated closely with the fraction of mature monocytes or dendritic cells in total bone marrow (Figure 5c). Highly aberrant progenitor cells gave rise to few mature myeloid cells. By contrast, progenitors more closely resembling healthy counterparts produced large numbers of mature myeloid cells. An AML with an NPM1 mutation (Figure 5c, blue and purple) produced more myeloid cells, compared to an NPM1-wt AML (Figure 5c, red and green) with a similar degree of aberration. AMLs with aberrant karyotype and CNVs (Figure 5c, red and blue) produced very few mature myeloid cells. Evidently, in bone marrow aspirates with low overall differentiation, many monocytes derived from residual healthy progenitors, while in bone marrow aspirates with many monocytes, these monocytes were derived from leukemic progenitors (Figure 5d). Leukemia-derived monocytes were enriched in a cell state with higher expression of MHC-II and several interferon response genes (IFITM1, IFITM3) (Figure 5e, Supplementary Table 2).

Together, these results suggest that leukemia-derived monocytes originate from incomplete differentiation blocks at the progenitor level and can mature along normal differentiation pathways. The stage of the differentiation block at an MPP, LMPP or promyelocyte stage determines the first-line chemotherapy response of an AML, and is subject to evolutionary selection. By contrast, the strength of the differentiation block is also encoded at the progenitor level and determines the degree of monocytic differentiation; however, this property is independent of the stage of the block.

### CloneTracer enables the discovery of leukemia specific markers in a differentiation-state aware manner

Our results indicated that leukemic stem cells, as well as leukemia derived monocytes, are rather healthy-like and thus particularly difficult to distinguish from their healthy counterparts. This raises the question if there are specific markers that can be used to identify and/or target healthy vs. leukemic cells of various differentiation stages, including stem cells and monocytes; such markers can possibly be identified by comparing between healthy and leukemic cells from the same patient, thereby avoiding batch effects, genetic background and other variables typically confounding healthy-cancer comparisons. To investigate this idea, we first used cohort B, since a larger number of surface markers and cells per patient was covered. We asked if there are markers that are overexpressed or depleted in leukemic cells of various differentiation stages, compared to healthy cells of the same stage. In 9 out of 16 combinations of patients, cell states and sampling timepoints available for comparison, CD11c was significantly overexpressed by leukemic cells, whereas CD49f was significantly enriched on healthy cells (Figure S7a). Thus, after accounting for differentiation stage specific expression changes (Figure S7b) by pre-gating with the monocyte marker CD14, an enrichment of healthy cells was obtained in the CD11c-CD49f+ gates, compared to the CD11c+CD49f-gates and vice versa (Figure 6a,b). We confirmed the specificity of the CD11c/CD49f marker combination by FACS sorting followed by Fluorescent In Situ Hybridization (FISH) in patient B.4. Leukemic cells carrying the monosomy 7 were enriched in the CD11c+CD49f-fraction, whereas healthy cells diploid for chromosome 7 were enriched in the CD11c-CD49f+ fraction (Figure 6c, S7c) (p=1×10^-10^). Similar enrichments were demonstrated in an independent patient with trisomy 8 (Figure S7d).

**Figure 6.**
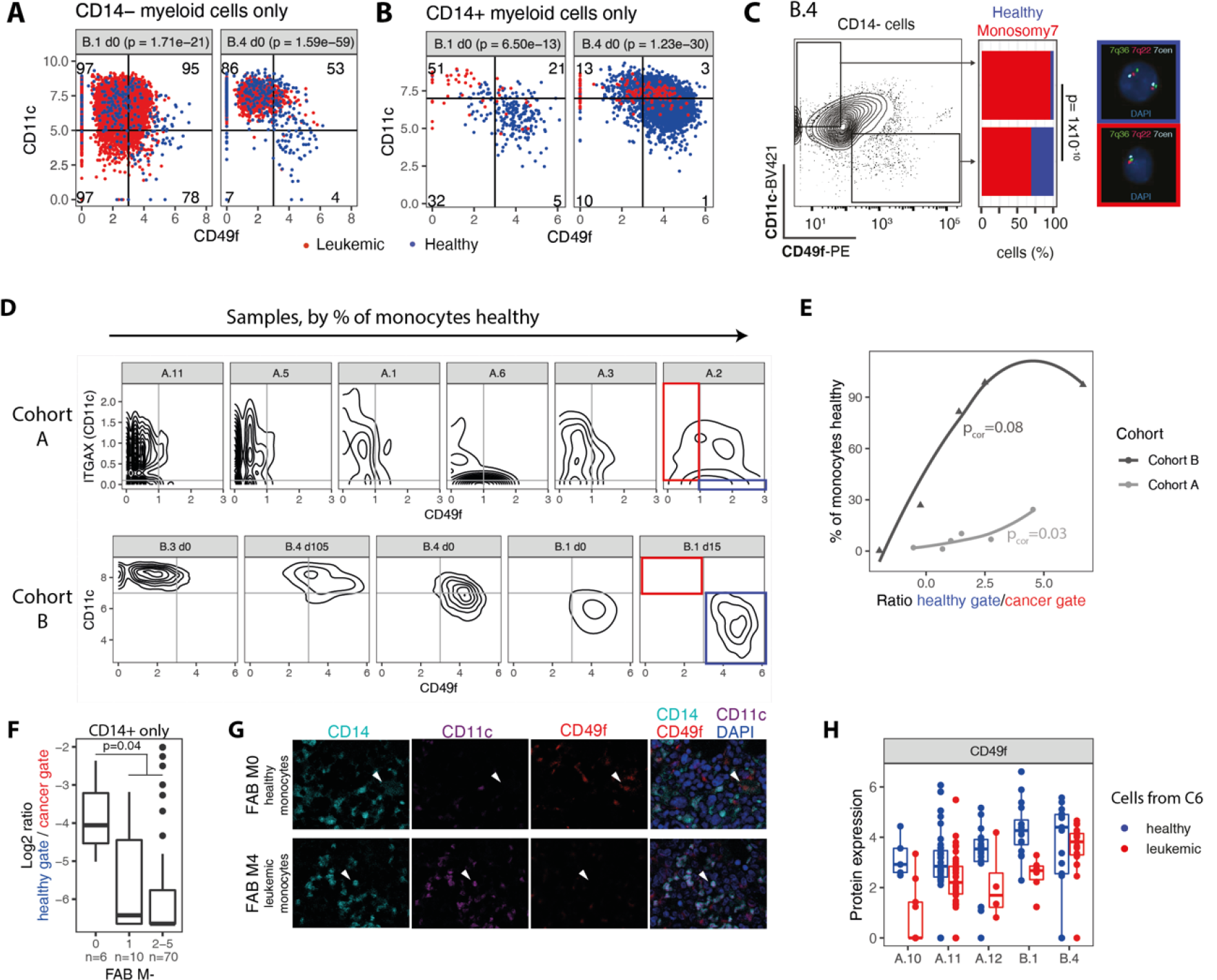
Discovery of leukemia specific markers by intra-patient differential expression testing. **a.** Scatter plots of the expression of CD49f and CD11c highlighting the clonal identity for patients B.1 and B.4. Only CD14-cells are shown. Numbers indicate the percentage of leukemic cells in each of the four quadrants. p values are from a Fisher test for the association of quadrant with clonal identity. **b.** Like a, except that only CD14+ cells are shown. **c.** FACS plot depicting CD11c and CD49f expression in pre-gated CD14-bone marrow cells of patient B.4. Bar Plots depict the fraction of healthy (blue) and leukemic (red) cells in sorted CD14-CD11c+CD49f- (n = 162 cells) and CD14-CD11c-CD49f+ (n = 151 cells) populations. p values were calculated using a Fisher test for the association of the sorting gates with clonal identity. Representative images show hybridization of three FISH probes for the chromosome regions 7 centromere (7cen, cyan), 7q22 (red) and 7q36 (green) in healthy (top/blue; two signals for each probe) and leukemic (bottom/red; one signal for each probe) cells. Nuclei were counterstained with DAPI. **d.** Density plots depicting the expression of ITGAX/CD11c and CD49f in monocytes across patients with at least 25 monocytes represented, and confident CloneTracer assignments. Individual panels are arranged by % of healthy monocytes. **e.** Scatter plot relating the fraction of healthy monocytes to the ratio between the gates indicated in red and blue in panel d. **f.** Tissue microarray data from n=86 patients. Ratio between the healthy and cancer gate (see panel d) was calculated as a function of FAB classification. FAB M0: undifferentiated; FAB M1: minimal differentiation; FAB M2-M5: varying differentiation. p value was calculated using a Wilcoxon test. **g.** Representative mid-optical sections of an CD14, CD11c and CD49f stained tissue micro array used for quantification in panel f (see also Figure S7e,f). Nuclei were counterstained with DAPI. Upper row: Representative fluorescence images of a FAB M0 classified AML patient. Arrows point to a potentially healthy CD14+/CD11c-/CD49f+ cell. Lower row: Representative images of a FAB M4 classified AML patient. Arrows point to a potentially leukemic CD14+/CD11c+/CD49f- cell. **h.** Box plot comparing the expression of CD49f in healthy and leukemic cells from cluster C6, see also Figure 4g.

To follow up on these discoveries, we integrated data from cohort A (where only CD49f had been covered at the protein level, and imputation with MAGIC^35^ was used to estimate CD11c expression levels from *ITGAX* RNA levels). Further, we analyzed tissue microarray data from 86 AML patients for expression of CD14, CD34, CD11c and CD49f (Figure S7e,f), as well as flow cytometry data of 87 AML patients for expression of CD14, CD34 and CD11c. Together, these results allow us to provide a perspective on the significance of CD11c and CD49f, and their significance as healthy vs. leukemia markers.

At the monocyte level, an increased presence of leukemia-derived cells was associated with a larger fraction of CD11c+CD49f- (Figure 6d,e) in both single cell cohorts. In the TMA data, we observed that in leukemias with differentiation (FAB M2-M5), putative leukemic CD14+ cells predominantly were CD11c+CD49f-. In undifferentiated leukemia (M0), residual, presumably healthy CD14+ cells showed a CD11c-CD49f+ phenotype (Figure 6f,g, S7e,f). In the flow cytometry data, we observed that an increased rate of monocytic differentiation of the AML coincided with an increased expression of CD11c on the mature monocytes (Figure S7g). Of note, this association was observed in all AML subtypes except those with inv(16) or TP53 complex karyotype AML. The CD11c+CD49f-state corresponded to the MHC-II state described above (Figure S7h, 5e). Thus, at the level of monocytes, CD11c and CD49f form a robust combination of markers to quantitate the fraction of leukemia content. This might be particularly helpful in AML diagnosis if large number of monocytes are present.

In stem cells, the CD11c+/CD49f- combination was informative in a subset of patients. In four out of five patients with healthy and leukemic cells in cluster 6, CD49f was more highly expressed by healthy cells from this cluster, i.e. by healthy dormant stem cells, in line with previous work on this marker^36^ (Figure 6h). The expression of CD11c on CD34+ cells was variable across and also within genotypes (Figure S7i,j). Accordingly, data integration by single cell transcriptomics was overall superior in identifying the multipotent leukemia stem cell cluster, as well as stem-like progenitor cells, compared to flow cytometry.

## Discussion

To investigate routes of cellular differentiation in AML, we have here introduced CloneTracer, a computational method for adding clonal resolution and identifying leukemic and healthy cells in single-cell RNA-seq data. CloneTracer accounts for the noise properties of nuclear SNV calls, mitochondrial SNV calls and CNV inference, integrates bulk data if available, and does not require prior knowledge of the clonal hierarchy. Thereby, CloneTracer extends upon DNA-seq specific error models^19–21, 37^ and upon models that require prior knowledge of the clonal hierarchy^22^. In the AML context, CloneTracer confidently identified healthy and leukemic cells in 13/19 patients. Certain types of chromosomal aberrations, such as inversions and translocations, can only be mapped at the single cell level with highly specialized, low-throughput protocols^38^. Furthermore, in droplet-based scRNA-seq protocols, SNVs in lowly expressed genes such as TET2 are difficult to amplify. For DNMT3a, coverage was obtained in approximately 20% of CD34+ cells; hence, pre-leukemic cells were difficult to distinguish from healthy cells with high confidence, unless they were marked with a mitochondrial mutation. Example data from healthy elderly individuals suggest that about 40% of hematopoietic cells should carry a mitochondrial clonal marker^39^, however, the variability of this phenomenon across larger cohorts has not been investigated. Although our method is therefore limited in distinguishing healthy and pre-leukemic cells, analyses using various types of biological validation (Figures 3, 6a-c) demonstrated low false positive and false negative rates in identifying fully leukemic cells, justifying the use of CloneTracer for the analysis performed here.

The availability of clonal information enabled us to clarify routes of leukemic differentiation. Of note, through the integration of data from all patients, we identified a cluster of stem cells that consisted of dormant, predominantly healthy stem cells and exclusively leukemic, active SCs with erythroid potential and a gene expression signature resembling label-retaining cells in xenotransplants^31^. Since both dHSCs/dLSCs and aLSCs had relatively consistent gene expression signatures across patients and healthy individuals, data integration of scRNAseq datasets represents a robust strategy for their identification. By contrast, these cells are difficult to enrich through flow sorting strategies: although dormant stem cells are CD34+CD38-, the large degree of inter-patient heterogeneity at the level of progenitors makes it difficult to develop universal purification schemes.

Downstream of LSCs, we observed a highly heterogeneous and aberrant compartment of immature myeloid cells. The patient specific stage of the differentiation arrest observed in this compartment determined the initial chemotherapy response of the patient. The strength of the block independently determined the overall degree of monocytic differentiation. Unlike cellular hierarchies identified from bulk data^40^, the availability of single cell resolution allowed us to distinguish stem cells from immature, “stem-like” progenitors, and pin-point a poor first-line chemotherapy response to the latter. This hypothesis goes in line with the kinetics of hematopoietic differentiation. Stem cells divide a few times per year, i.e. too slowly to play a role in determining immediate chemotherapy response, whereas progenitors divide up to several times per day^41^. By contrast, relapse in AML occurs at a time scale compatible with the activation of dormant stem cells. Therapy resistance and relapse therefore possibly originate from distinct cellular compartments.

Figure 7 summarizes our model of leukemic differentiation pathways, and suggests overall similarities to healthy hematopoietic differentiation, but with an aberrant myeloid progenitor compartment. Our model suggests that leukemic mutations, while present in stem cells and monocytes, mostly exert their effect in the epigenetic context of progenitors. This implies that AML evolution requires mutations in slowly dividing stem cells, although selection occurs at the level of progenitors. Such a model would be in line with the low number of genetic aberrations observed in most AMLs. Of note, our data is static, and cannot exclude the possibility that progenitor cells in AML might also de-differentiate to give rise to stem cells.

**Figure 7.**
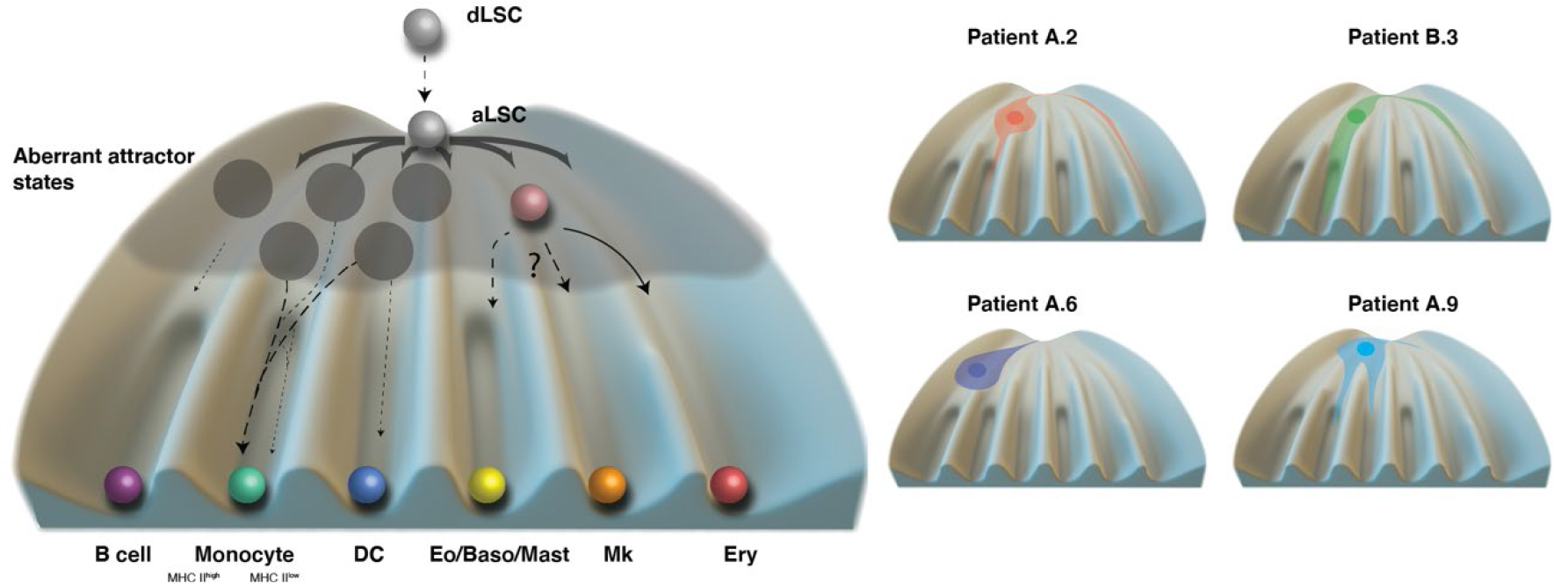
Model of leukemia differentiation. Adapted from a model of healthy hematopoietic differentiation^1^.

In sum, our data carry implications for the future development of therapeutic strategies: Our results indicate that in most if not all AML patients, there is a stem cell compartment distinct from the most immature progenitor cells. Hence, targeted therapies aimed at immature progenitors may increase the initial therapeutic response, but unless these therapies also target the actual stem cell compartment, the effect on relapse and long-term survival will be minimal.

## Acknowledgements

We thank Malte Paulsen, Beata Ramasz and Diana Ordonez for assistance with FACS sorting. Fengbiao Zhou, Anna Mathioudaki and Judith Zaugg for discussions. Laleh Haghverdi and Valerie Marot-Laussazaie for providing feedback on the mathematical model. We thank all members of GeneCore (EMBL) for assistance with the CITEseq experiments, as well as the DKFZ Single-Cell Open Lab (scOpenLab) for assistance with the MutaSeq/SmartSeq2 experiment. We thank the NCT CLB, Sektion Cell Biobanking, especially PD Dr. Katharina Kriegsmann, for processing and providing bone marrow samples. Figure 1a was created with BioRender.com. This work was financially supported by the German Bundesministerium für Bildung und Forschung (BMBF) through the Juniorverbund in der Systemmedizin ‘LeukoSyStem’ (FKZ 01ZX1911D to LV and SR), as well as the Verbundprojekt SMART-CARE (031L0212A to CMT), the Emerson foundation grant 643577 (to LV), grant PID2019-108082GA-I00 by the Spanish Ministry of Science, Innovation and Universities (MCIU/AEI/FEDER, UE), the German Research Foundation (DFG; Projects MU1328/18-1 and MU1328/21-1 and MU1328/23-1 to CMT) and the German Cancer Aid (DKH; Project 70113908 to CMT). LV acknowledges support of the Spanish Ministry of Science and Innovation to the EMBL partnership, the Centro de Excelencia Severo Ochoa and the CERCA Programme / Generalitat de Catalunya. AKL and JAK gratefully acknowledge the data storage service SDS@hd supported by the Ministry of Science, Research and the Arts Baden-Württemberg (MWK) and the German Research Foundation (DFG) through grant INST 35/1314-1 FUGG and INST 35/1503-1 FUGG. JAK acknowledges support of the Deutsche Gesellschaft für Hämatologie und Medizinische Onkologie e.V. (DGHO) and Deutsche José Carreras Leukämie-Stiftung e.V. through the José Carreras-DGHO-Promotionsstipendium.

## Author contributions

AKL, SR, CMT and LV conceived the project. CST, AKL, JAK, MJ, MA, AW and MB generated data and developed laboratory protocols. SBC, JAK, AKL, MB, PP and LV analyzed the data with support from MS and CR. SBC and LV developed the statistical model. JL and VB generated and processed raw sequencing data and supported data analyses. SB advised on antibody-oligo conjugation. AJ and MB performed and analyzed the FISH. LV and CMT supervised research. All other authors were involved in the acquisition and characterization of clinical specimen. LV, AKL, SBC and CMT wrote the manuscript and generated the figures. All authors have read and commented on the manuscript.

## STAR Methods

### Patients

Bone marrow samples from AML patients were obtained at the Heidelberg University Hospital after informed written consent using ethic application number S-169/2017. For demographic characteristics on sample donors, see Supplementary Table 3. Bone marrow aspirates were collected from iliac crest. Mononuclear cells were isolated by Ficoll (GE Healthcare, Chicago, Illinois, USA) density gradient centrifugation and stored in liquid nitrogen until further use. All experiments involving human samples were approved by the ethics committee of the University Hospital Heidelberg and were in accordance with the Declaration of Helsinki.

### Panel, exome, and bulk ATAC sequencing

For bone marrow samples form cohort A, CD3- and CD3+ cells were sorted by FACS and subjected to exome sequencing as described before^25^. GATK best practices were followed. Mutect2 with Tumor with match normal option was used for the identification nuclear mutations specific for each patient. We considered CD3- cells as tumor and CD3+ as normal.

Additionally, samples A.1, A.3, A.5, A.6, A.7, A.11, A.12, A.13, A.15 were subjected to bulk ATAC sequencing to identify mitochondrial mutations. Again, Mutect2 with Tumor with match normal option was used to identify variants in the mitochondrial genome.

Bone marrow samples from cohort B were sequenced at diagnosis time point with the Illumina TruSight Myeloid Sequencing Panel (Illumina, San Diego, California, USA) to determine the mutation status of leukemia driver mutations.

### Antibody-oligo conjugation

For markers where no commercial conjugates were available, azide-modified oligonucleotides were conjugated to purified antibodies (anti-human CD166, Clone 3A6 (Biolegend, 343902); anti-human GPR56, Clone 4C3 (Biolegend, 391902)) by the use of a DBCO-PEG5-NHS Ester (Santa Cruz Biotechnology, Dallas, USA) in a copper-free click reaction ^42^.

In brief, azide-containing storage buffer of purified antibodies was exchanged to PBS (pH 8.5) using the Amicon Ultra-0.5 NMWL 30 kDa Centrifugal Filter (EMD Millipore, Billerica, USA).

100 μg of PBS-buffered antibody was incubated with 2mM DBCO-PEG5-NHS in a final reaction volume of 100µL for 30 minutes at room temperature. The reaction was stopped by the addition of 100mM Tris HCl (pH 8) for 5 minutes at room temperature and non-reactive DBCO-PEG5-NHS was removed using the Amicon Ultra-0.5 NMWL 30 kDa Centrifugal Filter.

Azide-modified oligonucleotides were reconstituted in PBS before adding 30pmol per 1µg DBCO-functionalized antibody. The click reaction was conducted at 4°C for 18 hours. Unreacted oligonucleotides were removed using the Amicon Ultra-0.5 NMWL 50 kDa Centrifugal Filter and the final volume was adjusted to 100µL using PBS (pH 8.5). Conjugation products were confirmed on Ethidiumbromide (EtBr) stained 2% agarose gels, Coomassie brilliant blue (CBB) stained 4-12% polyacrylamide gels and by absorbance spectroscopy.

Azide-modified oligonucleotides were purchased from Biolegio (Biolegio, Nijmegen, Netherlands) and contained an antibody-specific barcode (bold), a PCR handle (italic) and a capture sequence (underlined). * indicates a phosphorothioated bond to prevent nuclease degradation:

CD166: 5’/Azide/CCTTGGCACCCGAGAATTCCA**CATTAACAGCGCCAA**<u>CAAAAAAAAAAAAAAAAAA</u> <u>AAAAAAAAAAAA*A*A</u>

GPR56: 5’/Azide/CCTTGGCACCCGAGAATTCCA**TCATATCCGTTGTCC**C<u>AAAAAAAAAAAAAAAAAAA</u> <u>AAAAAAAAAAA*A*A</u>

### CITEseq surface labeling, FACS sorting and GEM generation

Human bone marrow samples were thawed and stained using the CITEseq antibody pool (Supplementary table 1), as well as sorting antibodies according to the BioLegend protocol (https://www.biolegend.com/en-us/protocols/totalseq-a-antibodies-and-cell-hashing-with-10x-single-cell-3-reagent-kit-v3-3-1-protocol)

In cohort A, sorting was done using fluorophore-tagged antibodies from BioLegend (San Diego, USA) against human CD3 (clone UCHT; 1:30), CD34 (clone 581; 1:100), and GPR56 (clone CG4; 1:20). FACS sorting of live bone marrow cells was performed on a BD FACSAria equipped with a 70 µm nozzle to enrich for the following populations: CD3+, CD3-CD34+, CD3-CD34-, while aiming for a representation of 25%, 50%, 25%. When insufficient CD34+ were available, a maximum of CD34+ cells were sorted and GPR56 was used as a second sorting marker to enrich stem cells from the CD34- fraction. Population frequencies were recorded and accounted for in quantitative analysis of the single-cell RNA-seq data set. Sorted cells were loaded onto the Next GEM Chip G for a targeted cell recovery of 5000 cells following the manufacturer’s instructions (10x Genomics, CG000206 Rev D).

In cohort B, fluorophore-tagged antibodies against human CD34 (clone 581) and CD45 (clone HI30) (patients B.1, B.2, B.3), or CD56 (clone QA17A16) and CD45 (patient B.4) were used to enrich for following populations:

|  | B.1 |  | B.2 |  | B.3 |  | B.4 |  |  |
| --- | --- | --- | --- | --- | --- | --- | --- | --- | --- |
|  | d0 | d15 | d0 | d15 | d0 | d15 | d0 | d21 | d105 |
| total BM/CD45+ | 85% | 99.6% | 100% | 100% | 92% | 100% | 100% | 100% | 98% |
| CD45dim/CD34+ | 15% | 0.4% | 0% | 0% | 8% | 0% | - | - | - |
| CD45dim/CD56+ | - | - | - | - | - | - | 0% | 0% | 2% |

Population frequencies were recorded and accounted for in quantitative analysis of the single-cell RNA-seq data set. In cases where different biological samples were combined in the same GEM generation run, cells were labeled additionally with oligonucleotide coupled cell hashing antibodies (Biolegend, San Diego, USA). FACS sorting of live bone marrow cells was performed using DRAQ7 (1:1000; Biolegend, San Diego, USA) and Incucyte Caspase3/7 Red (1:5000; VWR International, Radnor, Pennsylvania, USA) on a BD FACSAria^TM^ Fusion equipped with a 100µm nozzle. Sorted cells were loaded onto the Next GEM Chip G for a targeted cell recovery of 8000 cells following the manufacturer’s instruction (10x genomics, CG000206 Rev D).

### Single-cell RNA sequencing

cDNA libraries were generated using the 10x genomics 3’ gene expression kit version 3.1 according to the manufacturer’s instructions (10x genomics, CG000206 Rev D). At the cDNA amplification step (step 2.2 of the 10x genomics protocol), additive primers for amplification of the ADT and HTO libraries were added according to the manufacturer’s instructions (Biolegend protocol: TotalSeq^TM^-A Antibodies and Cell hashing with 10x Single Cell 3’ Reagent Kit v3 or v3.1 (Single Index) Protocol, Step II). Following cDNA amplification (10x genomics protocol: step 2.3A), cDNA was split: 10 µL were used for generating Gene Expression (GEX) libraries and 5µL were used for generating antibody-derived tags (ADT) and hashtag oligo (HTO) libraries, respectively, according to manufacturer’s instructions (GEX: 10x genomics protocol: CG000206 Rev D, Step 3; ADT and HTO: Biolegend protocol: TotalSeq^TM^-A Antibodies and Cell hashing with 10X Single Cell 3’ Reagent Kit v3 or v3.1 (Single Index) Protocol, Step III). The remaining material was used to construct mitochondrial and targeted mutation libraries.

Final GEX, ADT and HTO libraries were quantified by Qubit and QC’ed on the Bioanalyzer.

Final GEX and ADT libraries were sequenced on separate lanes on a NovaSeq (Cohort A) or HiSeq4000 (Cohort B) with a targeted sequencing depth of 50,000-100,000 reads/cell (GEX) and 300 reads/antibody/cell (ADT), respectively. HTO libraries were sequenced with a targeted sequencing depth of 4000 reads/cell on a NextSeq500.

### Optimized 10x: Mitochondrial libraries

For the full-length amplification of mitochondrial cDNA, mitochondrial primers were pooled so that each mitochondrial primer is present at a final concentration of 0.9µM (mito primer mix). See Supplementary Table 4 for all primer sequences used in this protocol. 10 ng of amplified cDNA was added to a PCR master mix containing 50 µl 2X KAPA HiFi HotStart ReadyMix (Roche), 4 µl 10 uM Partial_Read1 primer, 2.5 µl mito primer mix, in a total volume of 100µL. PCR was run as follows: 1 cycle of 95C for 3 mins, 11 cycles of [98C for 20 secs, 67C for 1 min, 72C for 1 min], and 1 cycle of 72C for 5 mins followed by a 4C hold. PCR product was then cleaned with 1.5X (v/v) CleanPCR beads (CleanNA), followed by two washes of 80% ethanol, and eluted in 30 µl EB (Qiagen), after which it was quantified by Qubit and QC’ed by running 1-2 ng of DNA on an Agilent Bioanalyzer High Sensitivity chip. Sample bioanalyzer traces after this step are shown in Figure S1a.

Mitochondrial mutation libraries were then generated by tagmentation with an in-house produced wild-type transposase (Tn5)^43^. Briefly, transposome assembly and linker loading was carried out by adding 1 µl of 2 mg/ml Tn5 and 1 µl of annealed linker Tn5ME-B/Tn5MErev to 9 µl of water followed by incubation at 23C whilst shaking at 300 RPM for 30 mins. Assembled transposome was then diluted 1:100 with water. In our experience, it typically required four parallel tagmentation reactions to generate adequate yield for sequencing for a given sample. In a single tagmentation reaction, 1.5 ng of cDNA were added to 10 µl of diluted Tn5 and 10 µl of 4X tagmentation buffer (40 mM Tris-HCl, pH 7.4; 40 mM MgCl_2_), 10µL DMF for a total of 40 µL. Tagmentation reaction in the PCR was run as follows: 1 cycle of 55C for 3 mins, then a 10C hold. It is important that the PCR is already at 55C when the PCR tubes are placed in the instrument. After tagmentation, 10 µl of 0.2% SDS was added to the tagmented mixture and incubated at room temperature for 5 mins to neutralize the reaction. Once the transposase has been neutralized, the tagmented sample was added to a PCR master mix of 54 µl 2X KAPA HiFi HotStart ReadyMix, 6 µl of 100% DMSO (Thermo Scientific), 10 µl of 10 µM Targeted 10X primer, and 10 µl of 10 µM N7XX primer. This reaction mix was split into two PCR tubes and PCR was run as follows: 1 cycle of 72C for 3 mins, 1 cycle of 95C for 30 secs, 12 cycles of [98C for 20 secs, 60C for 15 secs, 72C for 30 secs], and 1 cycle of 72C for 3 mins followed by a 10C hold. After the PCR, all reactions were pooled and underwent two rounds of successive bead cleanup. In the first bead cleanup, 0.6X (v/v) CleanPCR beads were used, followed by two 80% ethanol washes and eluted in 50 µl EB. In the second cleanup, 0.6X (v/v) CleanPCR beads were again used, followed by two 80% ethanol washes and eluted in 15 µl EB. The final library was then quantified by Qubit and QC’ed on the Bioanalyzer. Representative bioanalyzer traces are shown in Figure S1b.

### Optimized 10x: Targeted genotyping libraries

Nuclear mutations were selected from panel or exome sequencing data by choosing non-synonymous variants in expressed genes, located < 1.5kb away from the end of the gene. Primers targeting mutations of interest were designed using a customized version of the TAPseq Bioconductor package^24^, https://github.com/veltenlab/CloneTracer/tree/master/primer_design

Four rounds of PCRs were then used to generate nuclear mutation libraries. The first three rounds of PCRs are gene-specific nested PCRs and sequencing adaptors and indices were added in the last PCR.

In the first round of PCR (PCR1), 10 ng of amplified cDNA from 10X was added to a PCR master mix containing 2.5 µl of pooled outer gene-specific primers (final concentration of each individual primer in the final pool 10 µM-100 µM), 5 µl of 1 µM Partial_Read1 primer, 20 µl of 5 M Betaine, 50 µl of 2X KAPA HiFi HotStart ReadyMix, and topped up to 100 µl with nuclease-free water. PCR with a heated lid was run as follows: 1 cycle of 95C for 3 mins, 11 cycles of [98C for 20 secs, 67C for 1 min, 72C for 1 min], and 1 cycle of 72C for 5 mins followed by a 4C hold. PCR product was then cleaned with 1.5X (v/v) CleanPCR beads (CleanNA), followed by two washes of 80% ethanol, and eluted in 15 µl EB (Qiagen). After each round of post-PCR cleanups, PCR products were quantified by Qubit and QC’ed by running 1-2 ng of DNA on an Agilent Bioanalyzer High Sensitivity chip. Example bioanalyzer traces are shown in Figure S1c.

In the second round of PCR (PCR2), 10 ng from PCR1 was added to a PCR master mix containing 2.5 µl of pooled middle gene-specific primers, 5 µl of 1 µM Partial_Read1 primer, 20 µl of 5 M Betaine, 50 µl of 2X KAPA HiFi HotStart ReadyMix (Roche), and topped up to 100 µl with nuclease-free water. PCR was run as follows: 1 cycle of 95C for 3 mins, 10 cycles of [98C for 20 secs, 67C for 1 min, 72C for 1 min], and 1 cycle of 72C for 5 mins followed by a 4C hold. Again, PCR product was cleaned with 1.5X (v/v) CleanPCR beads (CleanNA), followed by two washes of 80% ethanol, and eluted in 30 µl EB (Qiagen). Example bioanalyzer traces are shown in Figure S1c.

The third round of PCR (PCR3) was run separately for each target gene. 10 ng from PCR2 was added to a PCR master mix containing 2.5 µl of pooled staggered gene-specific primers (concentration of each primer in the final pool: 25 µM), 5 µl of 1 µM Partial_Read1 primer, 20 µl of 5 M Betaine, 50 µl of 2X KAPA HiFi HotStart ReadyMix (Roche), and topped up to 100 µl with nuclease-free water. PCR was run as follows: 1 cycle of 95C for 3 mins, 10 cycles of [98C for 20 secs, 67C for 1 min, 72C for 1 min], and 1 cycle of 72C for 5 mins followed by a 4C hold. PCR product was cleaned with 1.5X (v/v) CleanPCR beads (CleanNA), followed by two washes of 80% ethanol, and eluted in 30 µl EB (Qiagen).

Finally, libraries were uniquely indexed for each sample. To this end, 10 ng from PCR3 was added to a PCR master mix containing 2.5 µl of 10 µM SI primer, 2.5 µl of 10 µM RPI-N7XX primer (see supplementary table 1), 50 µl of 2X KAPA HiFi HotStart ReadyMix (Roche), and topped up to 100 µl with nuclease-free water. PCR was run as follows: 1 cycle of 95C for 3 mins, 8 cycles of [98C for 20 secs, 52C for 15 sec, 72C for 45 sec], and 1 cycle of 72C for 5 mins followed by a 4C hold. PCR product was cleaned with 1.5X (v/v) CleanPCR beads (CleanNA), followed by two washes of 80% ethanol, and eluted in 15 µl EB (Qiagen).

### Raw data processing

Gene expression data was processed using cellranger version 4.0.0 with default parameters for feature barcoding. Doublets were removed using scrublet^44^. For cohort A, cells with <1800 genes detected or >10% mitochondrial reads were removed. For cohort B, cells with <1000 genes detected or >40 % mitochondrial reads were removed. The data from the healthy reference individual (C.3) was downloaded from https://doi.org/10.6084/m9.figshare.13397987.v3 and not subjected to further quality filters.

Mitochondrial libraries were processed following the DropSeq standard workflow^45^ except that reads were aligned to the mitochondrial genome (GRCh38). Consensus mitochondrial reads were called using the fgbio tool CallMolecularConsensusReads (v. 1.3.0). Only reads from cell barcodes which were detected in the gene expression dataset were used for the downstream analysis. Nucleotide counts were extracted for each single cell using pysam (v. 0.15.3). The final output of the workflow is a list of single-cell matrices in which for each position of the mitochondrial genome the number of A,T,C and Gs UMIs are stored. Mitochondrial variants were identified as previously described^25^, and identified using bulk ATAC sequencing, where available (most of cohort A). The workflow was implemented in snakemake^46^ and can be found in https://github.com/veltenlab/CloneTracer/tree/master/library_processing/mitochondria

Nuclear SNV libraries were processed similarly to the mitochondrial libraries with the difference that reads were aligned to the complete human genome (GRCh38). Only reference and alternative alleles (identified by panel sequencing) were considered for the final count table. Due to the high number of PCR amplification steps, only UMIs supported by at least two reads were included in the analysis. The final count table contains the number of reference and alternative UMIs for each single cell and targeted mutation. The workflow to process nuclear SNVs libraries was written in snakemake and can be found in https://github.com/veltenlab/CloneTracer/tree/master/library_processing/nuclear-snv

### Analysis of single cell gene expression data

For projecting single cell data onto a reference atlas of healthy bone marrow, we used a workflow based on scmap^47^ as described^4^. Sample code for reference atlas projection is available at https://git.embl.de/triana/nrn/-/tree/master/Projection_Vignette. Thereby, we obtained uMAP coordinates, cell type labels, and myelocyte pseudotime, where applicable.

For unsupervised integration of all data sets, scanorama^48^ was used with default parameters to integrate across the three cohorts A, B and C, using the cohort as the batch. Scanorama components were then imported into Seurat and uMAP, nearest neighbor graphs, and clustering were computed using the default Seurat pipeline with default parameters^49^.

### Statistical modelling of clonal hierarchies and clonal assignments (CloneTracer)

See supplementary note for a full description of the CloneTracer model.

### Differential expression testing

For differential expression testing of surface antigens, we used Wilcoxon tests following library size normalization. For differential expression testing of RNA, we used MAST^50^. In all cases, comparisons were performed separately by patients, and the number of patients where the change was significant was used as an overall measure of significance and consistency.

### Fluorescent In Situ Hybridization

Human bone marrow samples were thawed and stained with fluorophore-tagged antibodies against CD45, CD3, CD49f, CD11c, CD14 and CD34 as described above (CITEseq surface labeling, FACS sorting and GEM generation). For antibody clones and titrated amounts, see Supplementary Table 1. Cells were collected by FACS sorting on a BD FACSAria^TM^, or BD FACSAria^TM^ Fusion, respectively, each equipped with a 100µm nozzle. Sorted cells were fixed on glass slides in methanol/acetic acid. Hybridization was performed according to the manufacturer’s instructions by using FISH probes for chromosome regions 6q21/8q24 and 7cen/7q22/7q36 (MetaSystems, Altlussheim, Germany), respectively. Interphase nuclei were validated using an automated scanning system (Applied Spectral Imaging, Edingen/Neckarhausen, Germany).

### Multispectral imaging of the bone marrow microenvironment using tissue microarrays

The frequency of different cell subsets in the bone marrow microenvironment in AML patients was analyzed by multispectral imaging (MSI). Formalin-fixed and paraffin embedded (FFPE), decalcified bone marrow samples were stained as described elsewhere^51^. The marker panel used for staining included antibodies directed against CD34, CD14, CD11c, CD49f. For antibody clones and dilutions, see Supplementary Table 1. All primary antibodies were incubated for 30 min. Tyramide signal amplification (TSA) visualization was performed using the Opal seven-color IHC kit containing fluorophores Opal 520, Opal 540, Opal 570, Opal 690 (Akoya Biosciences., Marlborough, MA, USA), and DAPI. Stained slides were imaged employing the PerkinElmer Vectra Polaris platform. To unify the spatial distribution analysis, × 20 MSI fields (1872 × 1404 pixel, 0.5 µm/pixel) were analyzed. Cell segmentation and phenotyping of the cell subpopulations were performed using the inForm software (PerkinElmer Inc., USA). The frequency of all immune cell populations analyzed and the cartographic coordinates of each stained cell type were obtained.

### Flow cytometry analysis of CD11c expression

Human BM samples were stained as described above and analyzed using the BD Symphony (see Supplementary Table 1 for a list of antibodies used). Cells were pre analyzed using FlowJo v10.8.1. Doublets and dead cells were excluded, as well as artefacts using a time gate. Remaining cells were exported using the channel values and imported into R. Further, the package Spectre v1.0.0 was used for batch correction, clustering, dimension reduction and visualization following the “discovery workflow with batch alignment using CytoNorm”. Using the summary table of the package, CD11c expression was investigated on CD34+ and CD14+ cell clusters.

### Data visualization

All plots were generated using the ggplot2 (v. 3.3.5), ComplexHeatmap (v. 2.6.2) and pheatmap (v. 1.0.12) packages in R 4.0.2 or FlowJo (v. 10.6.1, BD). Boxplots are defined as follows: the middle line corresponds to the median; the lower and upper hinges correspond to first and third quartiles, respectively; the upper whisker extends from the hinge to the largest value no further than 1.5X the inter-quartile range (or the distance between the first and third quartiles) from the hinge and the lower whisker extends from the hinge to the smallest value at most 1.5X the inter-quartile range of the hinge. Data beyond the end of the whiskers are called ‘outlying’ points and are plotted individually.

## Code availability

The implementation of the model and code for primer design is available at https://github.com/veltenlab/CloneTracer

## Data availability

Datasets including raw and integrated gene expression data, cell type annotation, metadata and dimensionality reduction are available as Seurat v3 objects through https://doi.org/10.6084/m9.figshare.20291628. Raw data will be made available in EGA.

## Supplemental information and legends

**Supplementary Methods and Supplementary Note.** Description of the CloneTracer model and its application to the AML patients profiled in this study.

**Supplementary Table 1.** Antibodies used for CITE-seq and flow cytometry.

**Supplementary Table 2.** Differential expression testing results.

**Supplementary Table 3.** Patient characteristics and bulk exome data.

**Supplementary Table 4.** Primer sequences.

---

**Figure S1.**
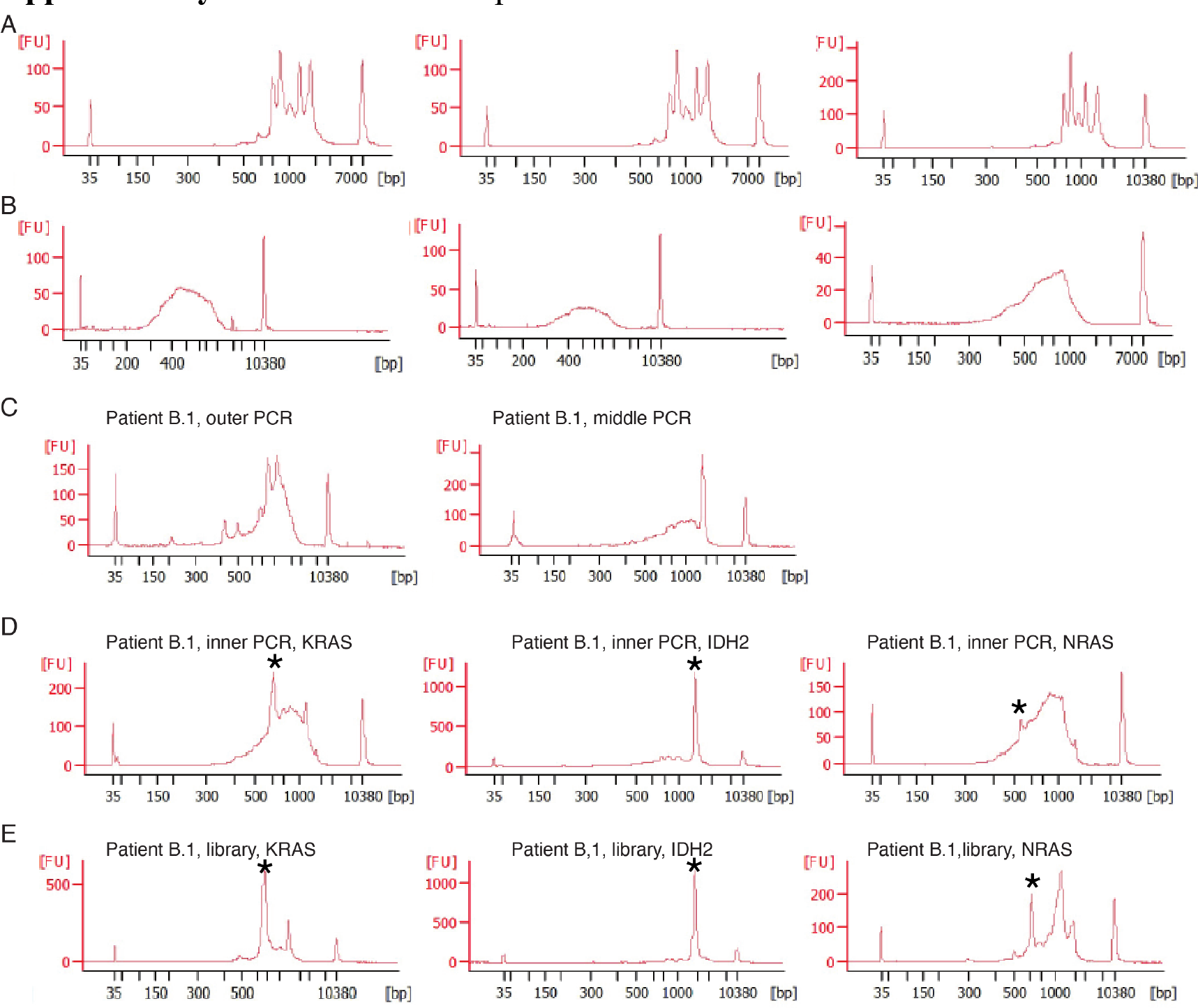
Bioanalyzer traces of “Optimized 10x” libraries. **a.** Mitochondrial library following PCR. Three representative libraries are shown. **b.** Final mitochondrial library following tagmentation and library PCR. Three representative libraries are shown. **c.** Nuclear mutation-targeting outer and middle PCRs, patient B.1. **d.** Inner PCR for KRAS, NRAS and IDH2 mutations in patient B.1. Asterisks indicate the expected product size. **e.** Final libraries for KRAS, NRAS and IDH2 mutations in patient B.1. Asterisks indicate the expected product size.

**Figure S2.**
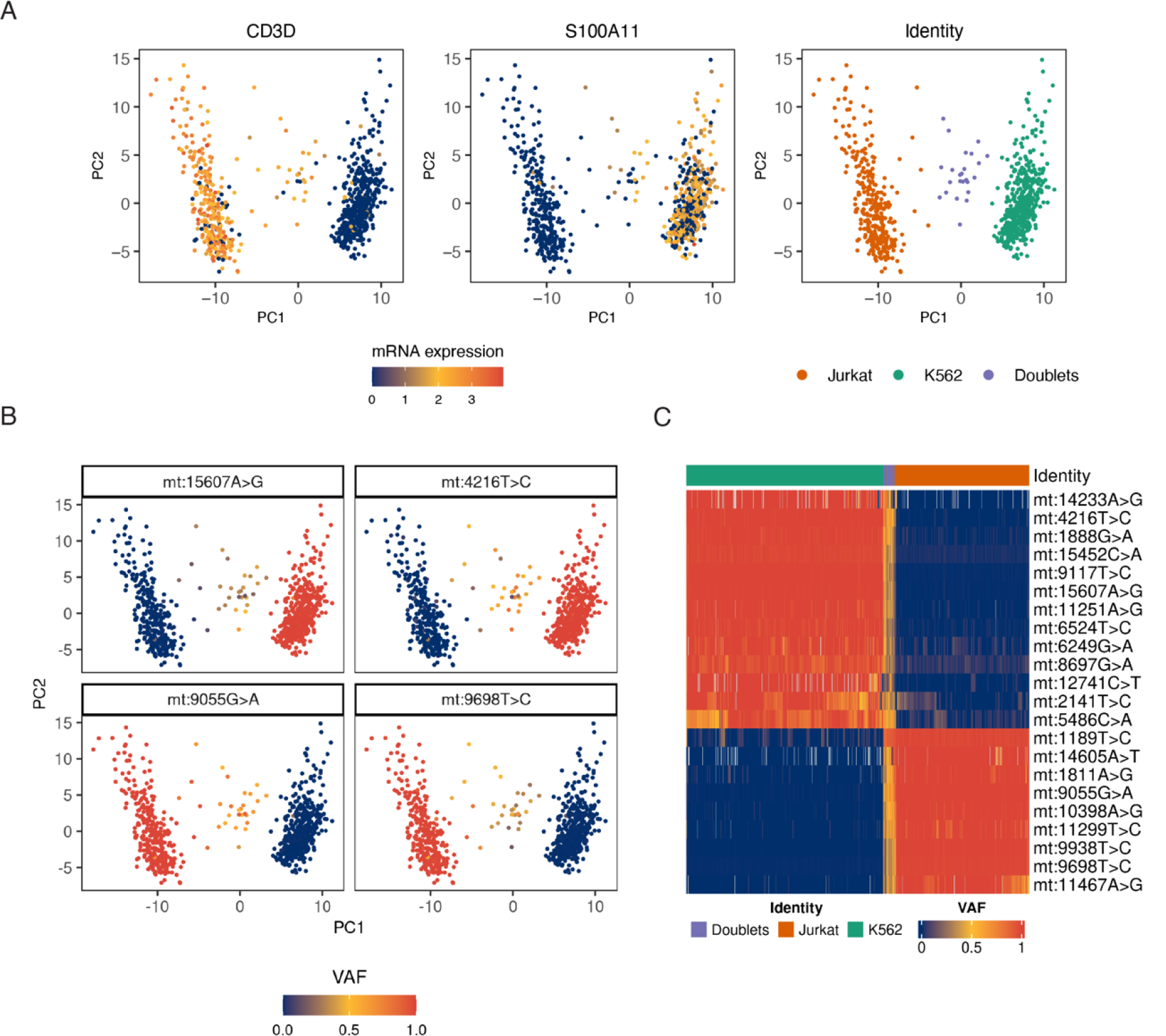
Cell line mixing experiment of Jurkat and K562 cells. **a.** Principal component analysis of gene expression. Score plots highlight the expression of a T cell gene (*CD3D*), a myeloid gene (*S100A11*) and assignment of cells as Jurkat, K562 or Doublets. Take note that doublets were removed from the main AML datasets using the scrublet algorithm prior to analysis. **b.** Score plots highlighting the variant allele frequencies of four mitochondrial variants that differ between Jurkat and K562. **c.** Heatmap depicting all mitochondrial mutations identified between these cell lines.

**Figure S3.**
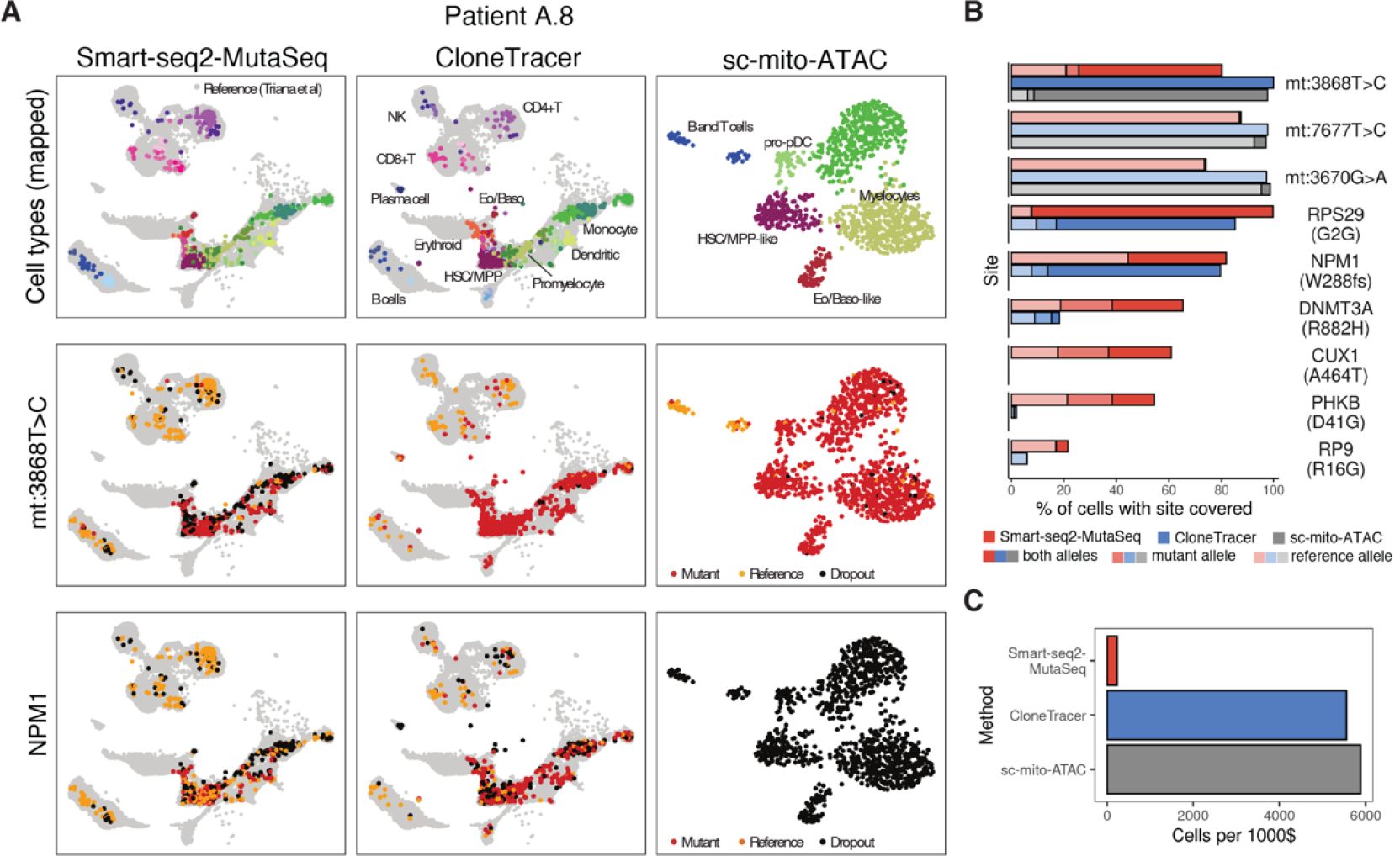
Comparison of the optimized 10x protocol to SmartSeq2/MutaSeq and sc-mito-ATAC. **a.** uMAPs illustrating the application of three single-cell genomic methods for clonal tracking to biological material from the same AML patient (A.8). Columns: Cells were profiled using MutaSeq (well-based RNA-seq), “Optimized 10x” (droplet-based RNA-seq) or sc-mito-ATAC seq (droplet-based ATAC-seq). Top row: Cells were mapped to a reference atlas^4^ using state of the art mapping algorithms^47, 52^, see also methods. Middle row: Mutational status for a mitochondrial mutation. Bottom row: Mutational status for a mutation in the gene NPM1. **b.** Coverage of nuclear and mitochondrial mutations identified in A.8. Nuclear mutations were identified by whole exome sequencing, mitochondrial mutations were identified using existing pipelines^25, 26^. Only myeloid cells are included. n=1142 (MutaSeq), 1072 (Optimized 10x), 1092 (sc-mito-ATAC) **c.** Bar chart comparing the throughput of the three methods.

**Figure S4.**
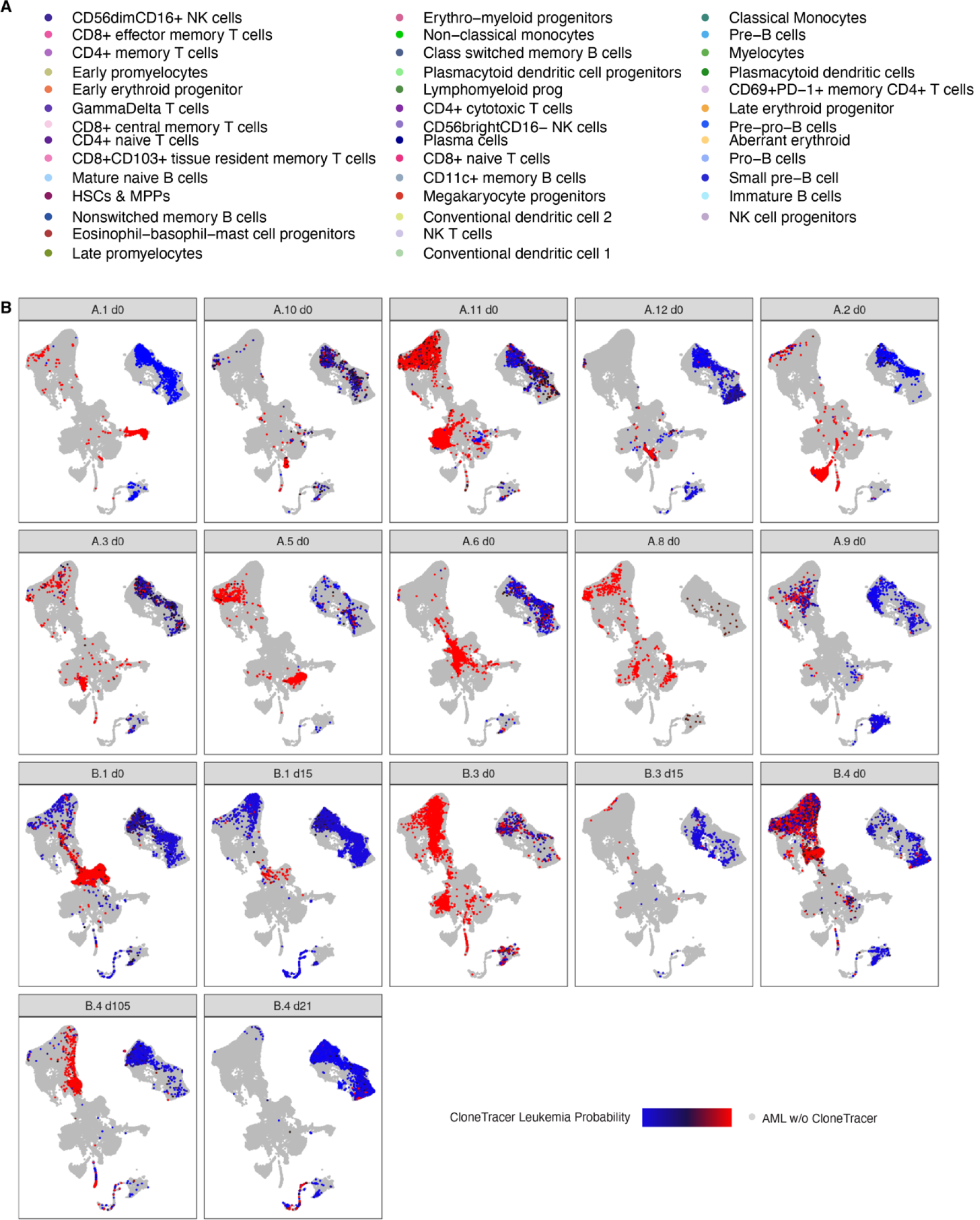
CloneTracer applied to two AML patient cohorts. See also main figure 2. **a.** Color legend for main figure 2c,d. **b.** CloneTracer assignments stratified by patient and time point.

**Figure S5.**
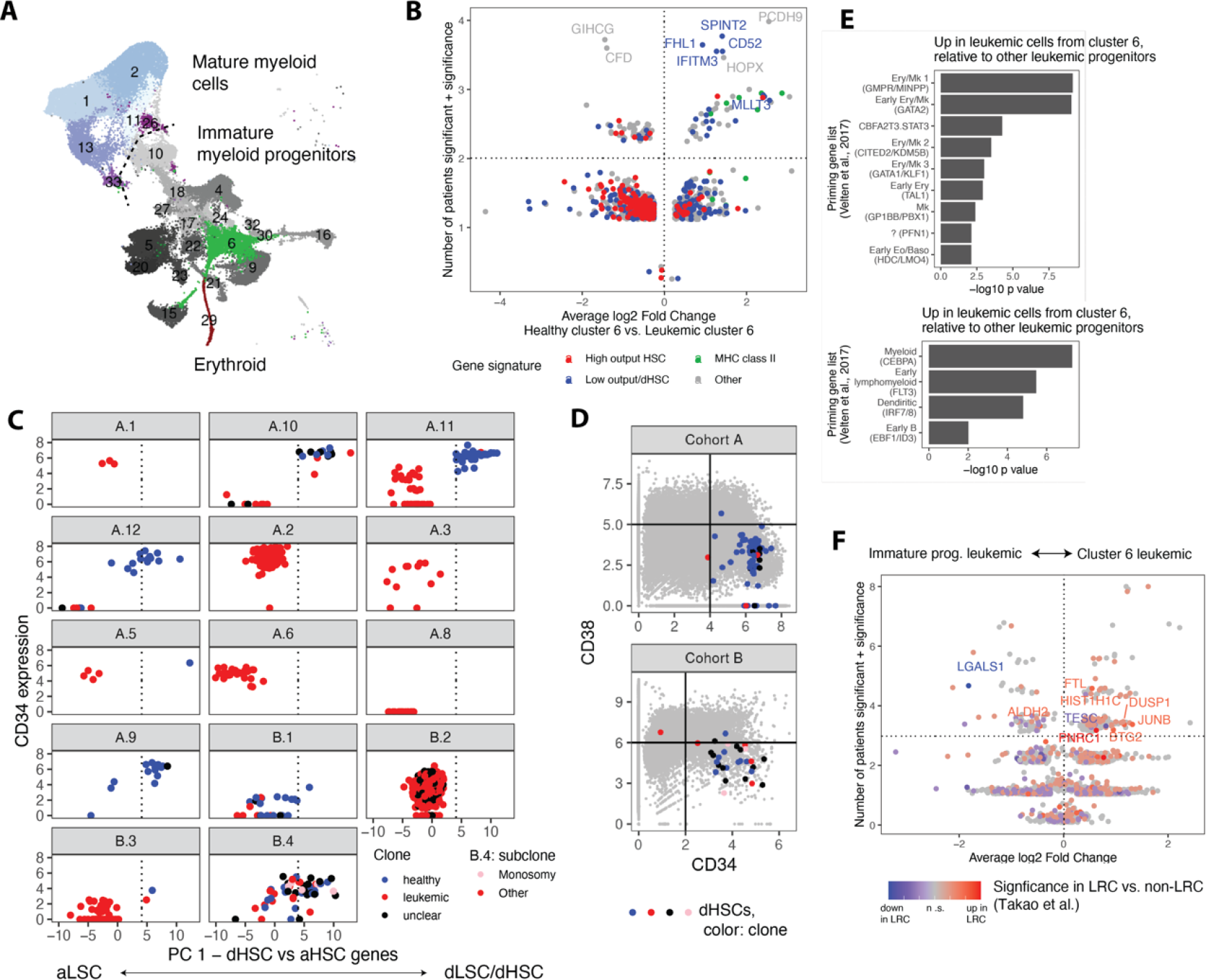
Analyses of healthy and leukemic stem cells, see also main figure 4. **a.** uMAP highlighting cluster identity. Clusters highlighted in shades of grey are referred to as “immature myeloid progenitors” in the main text, clusters highlighted in shades of blue as “mature myeloid cells”. **b.** Volcano plot as in main figure 4h, comparing healthy and leukemic cells from C6. n=5 patients were analyzed. Genes are colored by gene signatures for high output and low output HSCs identified by clonal tracking^34^ and dormant HSCs, identified by long-term label retention assays^33^. **c.** PCA of cells from cluster 6 performed jointly across all patients, but using exclusively genes from the dHSC signature^33^. Cells to the right of the dotted line were identified as putative dormant stem cells. **d.** Scatter plots depicting the surface marker expression of CD34 and CD38, highlighting the cells that were identified as putative dormant stem cells in panel c. **e.** Bar charts depicting the enrichment of gene signatures from ^1^ among genes significantly up- or down-regulated in at least 3 patients from figure 4i. **f.** Volcano plot as in main figure 4i, comparing leukemic cells from C6 to other leukemic immature myeloid cells from the same patient. n=13 patients with confident CloneTracer leukemia assignments were analyzed. Genes are colored using information on their expression in label retaining vs. non label retaining AML cells^31^.

**Figure S6.**
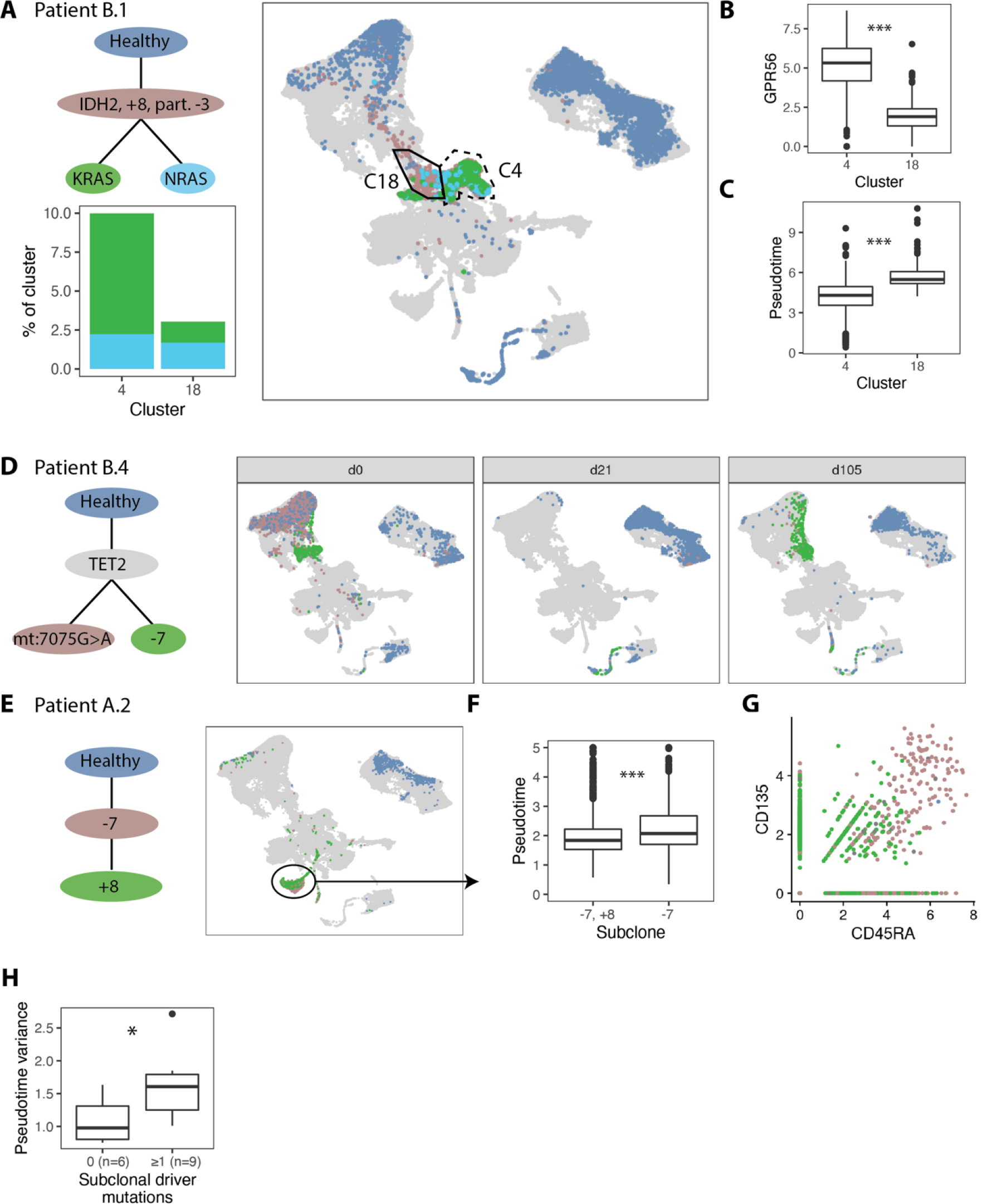
Analyses of subclones with CloneTracer. **a.** Top left: Clonal tree for patient B.1. Right: uMAP of day 0 data highlighting clonal identities. Brown dots also include cells with dropout of KRAS/NRAS. Two clusters (C4 and C18) are highlighted, see also Figure S5a. Bottom left: Representation of the KRAS and NRAS clones in clusters C4 and C18. **b.** GPR56 surface expression in clusters C4 and C18. **c.** Pseudotime on clusters 4 and 18. **d.** Clonal tree (left) and uMAP highlighting clonal identities (right) for patient B.4. **e.** Clonal tree (left) and uMAP highlighting clonal identities (right) for patient A.2. **f.** Boxplot of pseudotime within cluster 15 (highlighted), stratified by clone. **g.** Surface expression of CD45RA and CD135 for cells from cluster 15, color coded by clone. **h.** Box plot depicting the variance in pseudotime, stratified by the number of subclonal leukemic driver mutations identified from exome sequencing.

**Figure S7.**
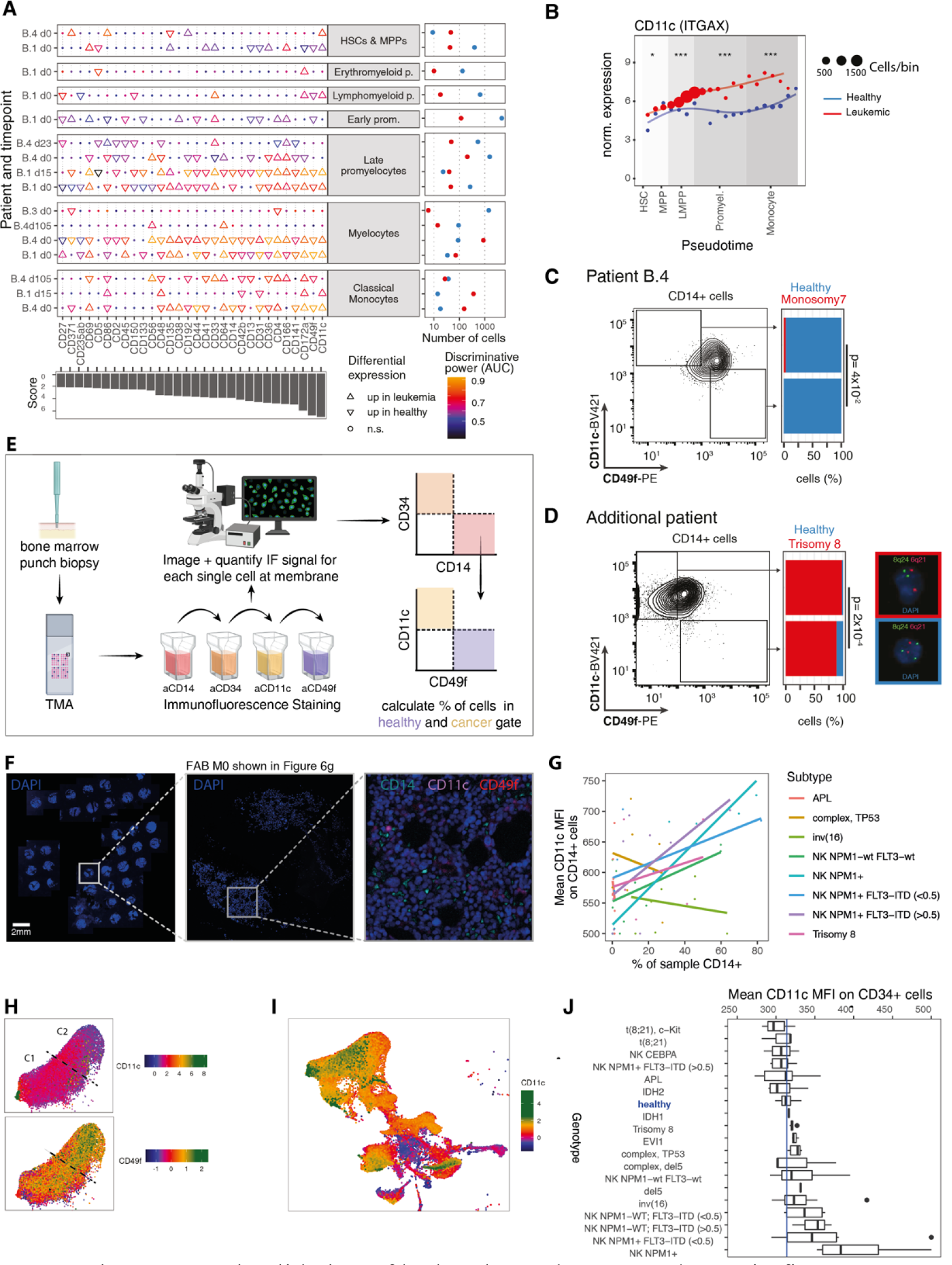
Discovery and validation of leukemia markers. See also main figure 6. **a.** Dot plot indicating intra-sample differential expression testing between healthy and leukemic cells. Upward triangles indicate surface markers with significant overexpression in leukemic cells, downward triangles indicate surface markers with significant overexpression in healthy cells. Color indicates discriminative power (area under the receiver operating characteristics curve). Right panel: Scatter plot indicating the number of healthy (blue) and leukemic (red) cells underlying these comparisons in each sample. Bottom panel: score evaluating the different markers, defined as the sum of AUC times the sign of the log fold change, if significant. **b.** Smoothened expression of CD11c over pseudotime of patient B.1 stratified by clone. Asterisks indicate significance of differential expression within the shaded area of pseudotime. ***: FDR < 0.001, **: FDR < 0.01, *: FDR < 0.1. p values are from a Wilcoxon test of library-size normalized ADT counts. Points indicate mean expression within 20 equally sized bins along pseudotime, point size indicates number of cells per bin. **c.** Quantification of healthy and leukemic cells in CD14+ cell fractions of patient B.4 by FACS and FISH. See main figure 6c for a detailed legend. **d.** Like c, but for an additional patient carrying a trisomy 8. **e.** Scheme illustrating the preparation and immunofluorescence staining of TMAs used for validating CD11c and CD49f as leukemia/healthy markers (see Figure 6f,g). **f.** Representative immunofluorescence images of a TMA (left) and a single punch biopsy of a FAB M0 classified AML patient (middle, right) used for validating CD11c and CD49f as leukemia/healthy markers (see Figure 6f,g). Nuclei were counterstained with DAPI. Scale bar 2mm. **g.** Scatter plot relating the CD11c mean fluorescent intensity on CD14+ cells to the fraction of bone marrow that is CD14+, and the genotype. Flow cytometry data from n=59 individuals is shown **h.** Top: Expression of CD11c on the uMAP of monocytes (cluster 1 and 2, n=13,067 cells). For Cohort A, MAGIC^35^ imputed expression of ITGAX is shown, for cohort B, surface protein expression is shown. Data was scaled to unit variance separately for both chorts. Bottom: Surface protein expression of CD49f on the uMAP. **i.** Expression of CD11c on the uMAP of myeloid cells. **j.** Bar chart relating the expression of CD11c on CD34+ cells to genotype across n=87 patients profiled by flow cytometry.

