## Supplementary Note for "Clonally resolved single-cell multi-omics identifies routes of cellular differentiation in acute myeloid leukemia"

### Supplementary information

#### Supplementary methods

##### *Statistical modelling of clonal hierarchies and clonal assignments*

The goal of our model is to (1) compute the evidence for any biologically reasonable clonal hierarchy, (2) identify the best clonal hierarchy and (3) compute the posterior probability (confidence) of the clonal assignment, see figures 1d and N1-8.

As input data to the model, we consider $M \in \mathbb{N}_0^{n\times m}$, a matrix of UMI counts supporting the mutant allele across $n$ single cells and $m$ mutations of interest, and $N \in \mathbb{N}_0^{n\times m}$, a matrix of total UMI counts falling on that site (reference or mutant). In the case of CNVs, $M$ contains the number of UMIs falling in the chromosomal region of interest and $N$ the total number of UMIs observed in the cell. A vector $t \in \{SNV, mtSNV, CNV\}^m$ specifies for each mutation whether it is a nuclear SNV, mitochondrial SNV or CNV. A vector u specifies for each cell the cell type identified by default single cell transcriptomic analysis. A vector v specifies for each cell the sample (e.g. timepoint) it is from when present.

As further input data, we consider two integer-valued matrices $a, b \in \mathbb{N}_0^{m\times s}$ that contain information from bulk experiments across s samples (e.g. timepoints), such as read counts supporting the mutant allele and total read counts from bulk exome or panel sequencing. When no such information is available, a and b are set to 0. Finally, $r \in \mathbb{R}_+^m$ specifies the known ratio in chromosome counts between affected and non-affected cells for CNVs (e.g. 1.5 for trisomy, 0.5 for monosomy, etc).

We assume that in reality, cells belong to a number $l \leq m+1$ of clones that are represented by a binary matrix $A$ of dimensions $m \times l$. For example, a clonal hierarchy where a healthy cells give rise to a founding clone that in turn gives rise to two independent sub-clones would be represented by

$$A = \begin{pmatrix} 1 & 1 & 1 & 0 \\ 1 & 0 & 0 & 0 \\ 0 & 1 & 0 & 0 \end{pmatrix} \qquad (1)$$

A vector $\pi \in \{1, \ldots, l\}^n$ maps every cell to a clone, i.e. a column in A. A list of simplices $\psi_s$ specifies the marginal probabilities of each clone in each sample (e.g. timepoint), i.e. the fraction of each clone in the population of cells.

Assuming that A and $\pi$ are known, data is generated through the following process, parametrized by a set of further parameters (FPR, c, h, g) defined below. For a graphical representation of the model, see Figure N1a.

If mutation j is a nuclear or mitochondrial SNV, we assume that mutant UMIs $M_{.j}$ are either false positives created from a background poisson process, or true positives sampled from the total number of UMIs $N_{.j}$ using a Beta-Binomial distribution. This modelling choice is justified in Figure N1b-c and full posterior predictive checks for all patients are provided in Figure N1d-f. The use of a Binomial sampling process (McCarthy et al., 2020) is inappropriate, as both mitochondrial and, more so, nuclear counts are overdispersed (Figure N1b), possibly as a consequence of allele-specific gene expression. Thus,

$$p(M_{ij}|A_{j,\pi(i)}, h_{t(j)}, FPR_{t(j)}, c_j) = \begin{cases} pPoisson(M_{ij}|FPR_{t(j)}) \text{ if } A_{j,\pi(i)} = 0 \\ pBetaBinom(M_{ij}|N_{ij}, h_{t(j)}c_j, (1-h_{t(j)})c_j) \text{ else} \end{cases} \quad (2)$$

Here, FPR is a false positive rate defined independently for each class of mutation, h is the heteroplasmy of the mutation and c is a concentration parameter specifying the degree of over-dispersion. For nuclear SNVs, we assume that within one cell 50% of DNA molecules are mutated. Due to this assumption, we set strongly informative prior on h (e.g. a Beta distribution with shapes 1000,1000), thereby avoiding non-identifiabilities in the case of nuclear SNVs with low coverage. Different priors can be used e.g. for loss-of-heterozygosity SNVs, or mutations that strongly affect RNA stability. The concentration parameter is mutation-specific as we observed that mutations with lower coverage (e.g. *DNMT3A*) tend to be more overdispersed than SNVs falling in highly expressed genes such as *NPM1*. By contrast, for mitochondrial SNVs, the model allows to infer the heteroplasmy individually for each mutation (i.e. fraction of mutated mitochondrial transcripts in a cell carrying the mutation) from the data, and a flat prior is used while the concentration parameter is shared across mtSNVs. For FPR and c, weakly informative priors are used, see source code for a full specification of priors.

If j is a CNV, we assume that the UMIs on the affected chromosomal region are drawn from the total number of UMIs in the cell using beta-binomial sampling.

$$p(M_{ij}|A_{j,\pi(i)}, h_j, g_j, c_{t(j)}) = \begin{cases} pBetaBin(M_{ij}|N_{ij}, g_j c_{t(j)}, (1-g_j)c_{t(j)}) \text{ if } A_{j,\pi(i)} = 1 \\ pBetaBin(M_{ij}|N_{ij}, h_j c_{t(j)}, (1-h_j)c_{t(j)}) \text{ if } A_{j,\pi(i)} = 0 \end{cases} \quad (3)$$

Importantly, the observed CNV ratio constrains the ratio between on-target reads in cells carrying the CNV vs. cells not carrying the CNV.

$$p(r_j|g_j, h_j) = pNorm(r_j|g_j/h_j, 0.05) \quad (4)$$

We observed that the fraction of reads on the affected chromosome depends on the cell type, e.g. the fraction of RNA molecules from chromosome 7 is much decreased in plasma cells compared to healthy cells, also in individuals not affected by the copy number variant (Figure N2a). We therefore additionally formulated a model accounting for this cell type covariate by replacing g and h by cell type specific parameters

$$p(M_{ij} | A_{j,\pi(i)}, h_{j,u(i)}, g_{j,u(i)}, c_{t(j)}) = \begin{cases} pBetaBin\left(M_{ij} | N_{ij}, g_{j,u(i)} c_{t(j)}, \left(1 - g_{j,u(i)}\right) c_{t(j)}\right) \text{ if } A_{j,\pi(i)} = 1 \\ pBetaBin\left(M_{ij} | N_{ij}, h_{j,u(i)} c_{t(j)}, \left(1 - h_{j,u(i)}\right) c_{t(j)}\right) \text{if } A_{j,\pi(i)} = 0 \end{cases} \quad (5)$$

For this more complex model, we inferred a cell type specific prior on the fraction of UMI on the affected chromosome from patients not affected by the CNV such that

$$logit(h_{j,u(i)}) \sim Norm\left(x_{u(i)}, \sigma\right) \quad (6)$$

Detail on the estimation of the prior (cell type specific average fraction of UMI on the affected chromosome $x_{u(i)}$ and its variation across patients $\sigma$) is provided in figure N2b,c. Posterior predictive checks confirm that this model describes the data better, compared to the simple, cell-type invariant model (Figure N2d). Application of the cell type specific model was only necessary in samples B.2 and B.4.

Finally, to include data from bulk measurements, allele frequencies are computed from clonal frequencies as follows

$$AF_j = 0.5 * \sum_{l \leq m} A_{jl} \psi_l \quad (7)$$

Then,

$$p\left(a_j | AF_j\right) = Binom\left(a_j | b_j, AF_j\right) \quad (8)$$

Together, equations 2 and 3 specify the likelihood of the single cell data, given the structure of the tree, clonal assignments, and all remaining parameters (Called $\theta$ in the following):

$$\mathcal{L}(M | A, \pi, \theta) = \prod_i \prod_j p(M_{ij} | N_{ij}, A_{j,\pi(i)}, \theta) \quad (9)$$

Equations 4 and 8 define the likelihood of additional observations available to the model, such that the final likelihood becomes

$$\mathcal{L}(M, a, r|A, \pi, \psi, \theta) = \mathcal{L}(M|A, \pi, \theta) * \prod_j p(a_j|A, \psi) * \prod_{j \in CNV} p(r_j|\theta) \quad (10)$$

Where CNV is the set of all CNV mutations.

We implemented the model in pyro, a probabilistic programming language that efficiently fits variational posterior distributions using stochastic variational inference with a highly parallel, GPU-based backend based on pytorch (Bingham et al., 2018). For a given tree A, we sample node attachment from the clonal fractions:

$$\pi_i \sim \text{Categorical}(\psi_{v(i)}) \quad (11)$$

and initially marginalize out the node attachment vector $\pi$. Priors on all other parameters of the likelihood functions (2-11) are specified above and in the source code, and are weakly informative unless specified above. We noticed that the parameters of our model (in particular, the clonal fraction $\psi$) strongly depend on the choice of A, making a joint inference of A and the remaining parameters inefficient.

We therefore perform model comparison with fixed A, using the evidence lower bound (ELBO). CloneTracer offers the possibility of generating all possible trees that are compatible with the three-gamete rule and the infinite site hypothesis and compute the ELBO for each configuration, however, in the case of larger numbers of mutations, this brute-force approach leads to unreasonably long runtimes and we therefore additionally implemented a search heuristic, where all tree configurations for the two most highly covered variants are explored and compared, and mutations are subsequently added to this tree in the order of their coverage.

We ensure convergence of the ELBO by plotting its value over iterations; in general, 400 iterations, starting with 5 particles, are sufficient for the ELBO to converge in patients harboring SNVs. For patients with CNVs 1000 iterations were selected to ensure convergence.

Once A is identified based on the ELBO, and all other parameters with the exception of the node attachment vector $\pi$ are inferred, we infer $\pi$ given the variational posteriors for all other parameters.

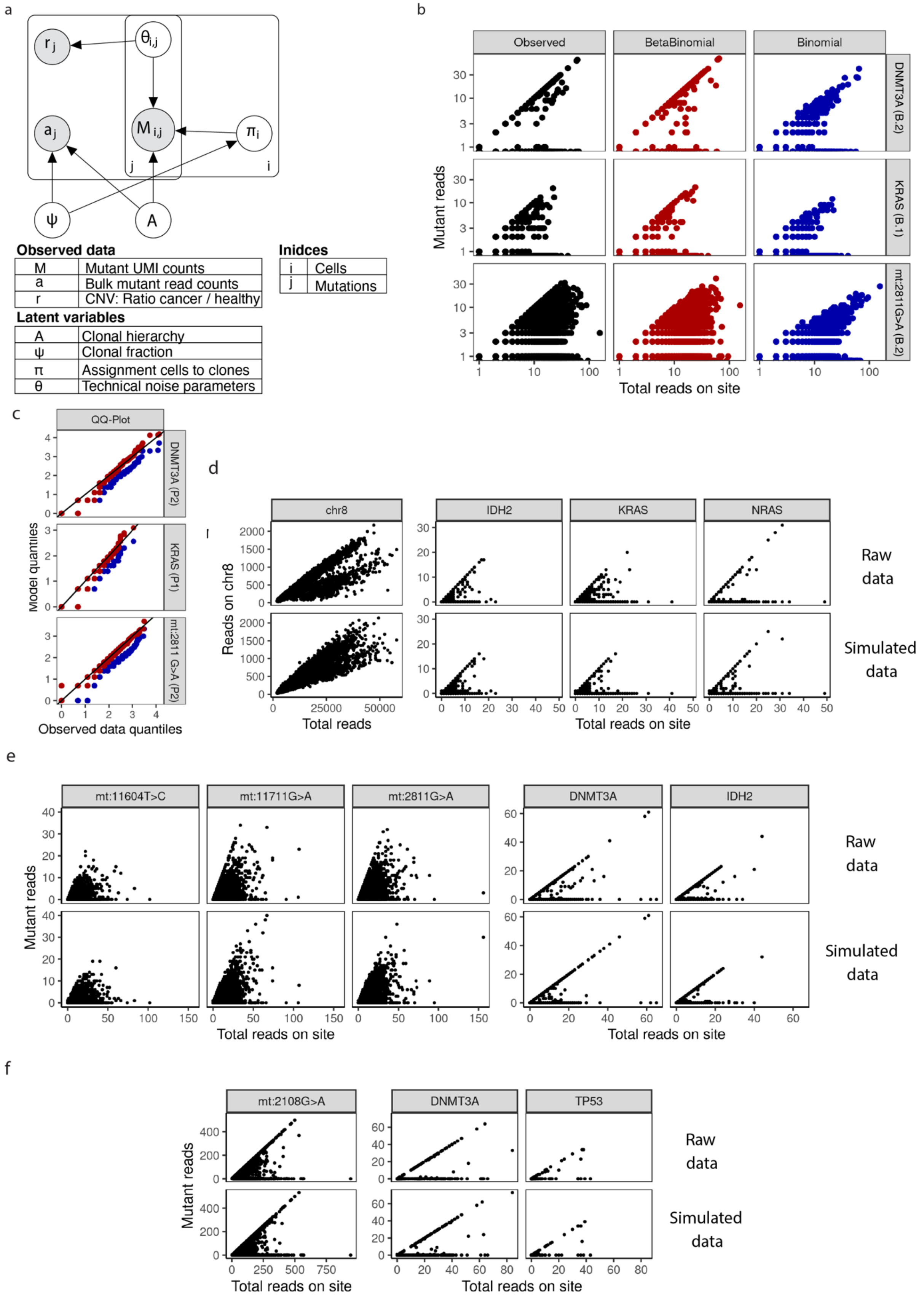
a
Observed data
M Mutant UMI counts
a Bulk mutant read counts
r CNV: Ratio cancer / healthy
Latent variables
A Clonal hierarchy
ψ Clonal fraction
π Assignment cells to clones
θ Technical noise parameters
Inidces
i Cells
j Mutations
b
Observed
BetaBinomial
Binomial
DNMT3A (B.2)
KRAS (B.1)
mt:2811G>A (B.2)
Mutant reads
Total reads on site
c
QQ-Plot
DNMT3A (P2)
KRAS (P1)
mt:2811 G>A (P2)
Model quantiles
Observed data quantiles
d
chr8
IDH2
KRAS
NRAS
Reads on chr8
Total reads
Total reads on site
Raw data
Simulated data
e
mt:11604T>C
mt:11711G>A
mt:2811G>A
DNMT3A
IDH2
Mutant reads
Total reads on site
Raw data
Simulated data
f
mt:2108G>A
DNMT3A
TP53
Mutant reads
Total reads on site
Raw data
Simulated data

**Figure N1.** Justification of basic modeling choices. **a.** Graphical display of the model, grey: observations, white: latent variables. **b.** Number of mutant reads plotted against number of total reads for three mutations (left column, black dots). The data was fitted using a mixture of a binomial sampling process to describe dropout of mutant reads in mutated cells and a poisson process to describe background false positive mutant reads in healthy cells (right column, blue dots). Alternatively, an overdispersed beta-binomial sampling process was used (central column, red dots). **c.** Quantile-quantile plots comparing the modeled beta-binomial and the binomial sampling process to the data. Red: Beta-Binomial model, blue: binomial model. **d.** Posterior predictive check for the full model. Top row: Raw data for patient B.1. Bottom row: The complete model described in the supplementary methods was fit to the data. Subsequently, mutant read counts were simulated from the model while keeping the total read counts and the clonal assignments fixed (posterior predictive check). Clonal assignments were kept fixed for this analysis since the model does not describe the relationship between total read count and cell type. **e.** Raw data and Posterior predictive check for patient B.2. **f** Raw data and Posterior predictive check for patient B.3.

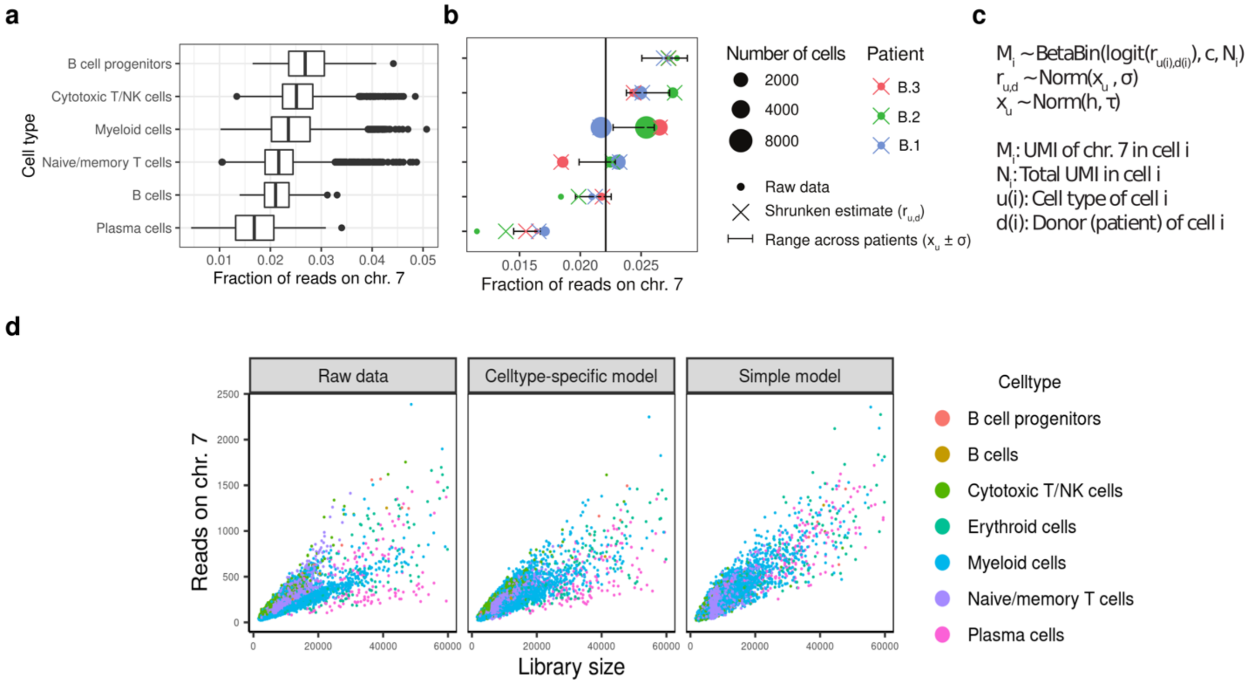

**Figure N2.** Accounting for cell type as a covariate in CNV analyses. **a.** Boxplot depicting the UMIs on chromosome 7 as a fraction of the total number of UMIs per cell. Data from patients B.1-B.3 (not affected by a monosomy 7) are shown. **b.** Inference of a cell type specific prior. Fraction of UMIs on chromosome 7 in different cell types is plotted; circles denote raw observed UMI fraction per patient and cell type, crosses denote a shrunken estimate, and the horizontal bar indicates the posterior estimate on the cell type specific UMI fraction and its standard deviation across patients. Inference was done using stan from the hierarchical model specified in panel c, and all cells from patients B.1, B.2 and B.3. **c.** Model used for estimating the fraction of UMIs on chromosome 7 per cell type, and its variance across patients. **d.** Raw data and Posterior predictive check for patient B.4, using both the simple model with no cell type covariate, and the cell type specific model.

#### Supplementary Note

##### *Application of CloneTracer to A and B cohorts of AML patient samples*

CloneTracer relies on well-covered clonal markers for the high-confidence identification of healthy and leukemic cells as well as sub-clones when present. Several types of genomic aberrations can be leveraged as clonal markers: nuclear single nucleotide variants (SNVs) affecting highly expressed genes (e.g. *NPM1*), mitochondrial SNVs (mtSNVs) and copy number variants (CNVs). For samples which only harbor nuclear SNVs occurring in lowly expressed genes, CloneTracer is unable to infer the clonal hierarchy due to the high dropout levels (see section *samples without well-covered clonal markers)*. As a result, healthy and leukemic cells cannot be identified confidently.

We selected nuclear mutations to include in this analysis as follows: All mutations with a distance of <1.5kb to the 3' end an average expression of >0.2 UMIs per cell in the patient were selected for primer design. Mitochondrial mutations were selected based on bulk ATAC sequencing, where available, or as previously described (Velten et al., 2021) (see also methods).

When it comes to selecting the clonal hierarchy for a particular sample, CloneTracer can output more than one possible tree configuration with equal statistical evidence. We often observe this behavior when poorly covered mutations are included (e.g. *DNMT3A*). In those cases, we always selected the mutation hierarchy with the lowest number of nodes. As a result of our maximum parsimony approach to select the clonal hierarchy, in samples where pre-leukemic mutations are present (e.g. mutations in *DNMT3A*), we cannot exclude that among cells identified as healthy, there are cells carrying pre-leukemic mutations. This is due to the low coverage of these type of mutations.

In the following subsections we describe in detail the output of CloneTracer when applied to AML samples from cohorts A and B.

##### *Samples with highly covered nuclear SNVs*

*NPM1 288fs* mutation is observed in around 30% of AML patients (Benard et al., 2021). Since the gene is highly expressed in immune cells, it can be leveraged as clonal marker (Figure N3). We often observe *NPM1* mutations co-occurring with *DNMT3A* SNVs in the same patient. In some cases, samples are made up of a unique clone such as patient A.9. In others, mtSNVs labelled subclones downstream of the initial *NPM1* founding clone (patients A.12, A.5 and A.8).

It is worth noting that synonymous mutations affecting highly expressed genes can also be used as clonal markers for CloneTracer. Patient A.8 is an example of this with a SNV in *RPS29*, a ribosomal gene which is covered in every single cell. *RPS29* mutation co-occurred with NPM1 mutation in this patient increasing the confidence in the distinction between healthy/pre-leukemic and leukemic cells. Only in this patient CloneTracer confidently selected a clonal hierarchy in which *DNMT3A* occurred before *NPM1* mutations. However, as mentioned in the section above due to the poor coverage of *DNMT3A* variants, the distinction between healthy and pre-leukemic cells is not possible.

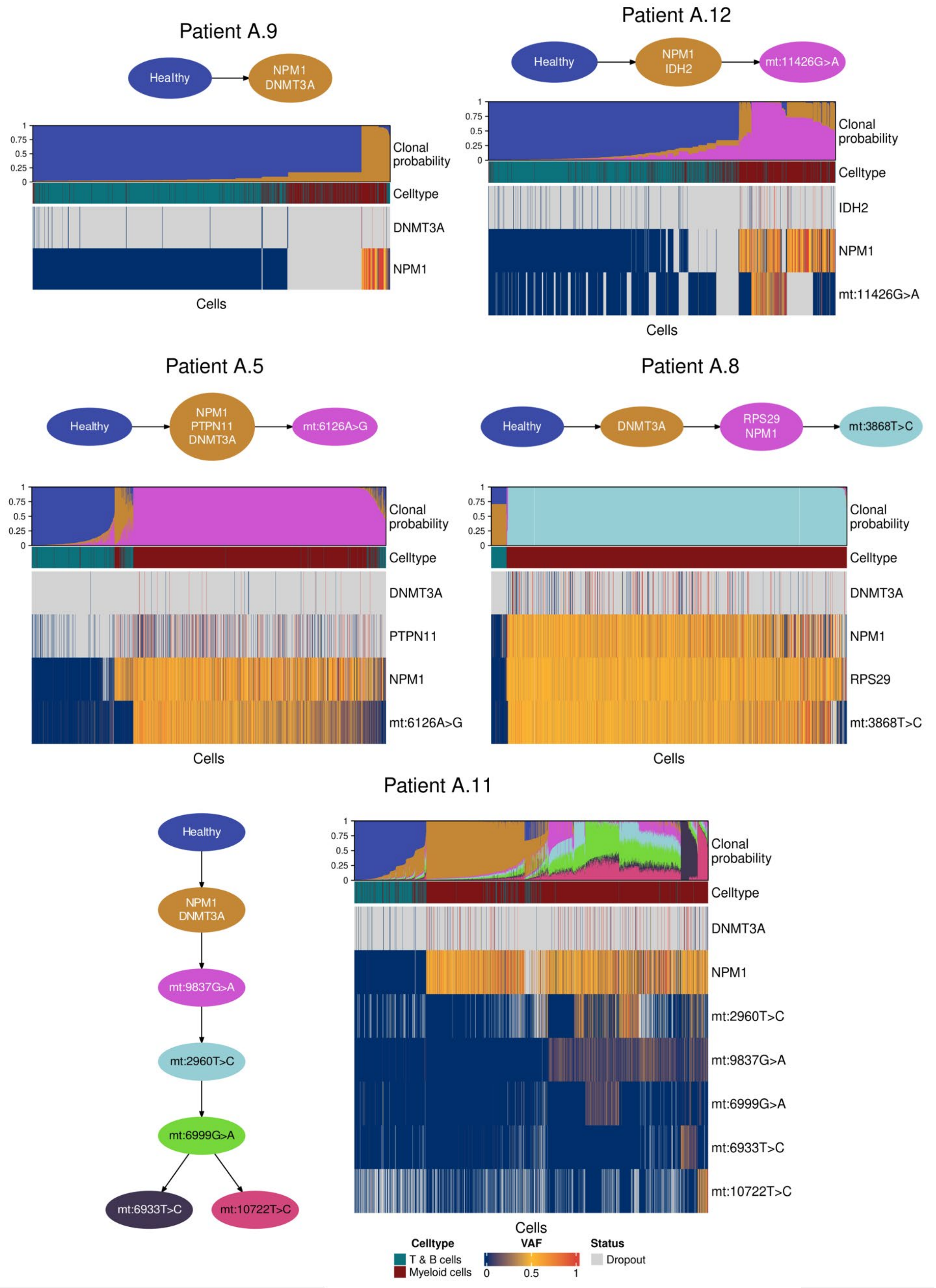

**Figure N3.** Application of CloneTracer to AML samples with well-covered nuclear SNVs. For each patient the clonal hierarchy inferred by the model is shown. The heatmaps display for each single cell the fraction of mutant reads (VAF, variant allele frequency) in all mutations as well as the cell type identity. Gray indicates that neither reference nor mutant reads were observed, termed as dropout. The clonal probabilities inferred by CloneTracer are shown at the top of the heatmap. Columns are ordered by decreasing clonal probability from the top to the bottom of the inferred clonal hierarchy.

In patient A.11, CloneTracer inferred a more complex hierarchy downstream of the *DNMT3A-NPM1* founding clone labelled by mitochondrial variants.

##### *Samples with mitochondrial SNVs as clonal markers*

Previous work has shown that mtSNVs can be used for lineage tracing in human samples (Lareau et al., 2020; Ludwig et al., 2019; Miller et al., 2022; Penter et al., 2021; Velten et al., 2021). In 3 patients, well-covered mitochondrial variants were present in the founding clone and therefore leveraged as clonal markers for the distinction of healthy and leukemic cells (Figure N4).

For patient B.3, CloneTracer inferred a clonal hierarchy with a single clonal population. This clone also harbors mutations in *DNMT3A* and *TP53* which are often found in individuals with clonal hematopoiesis, indicating that all leukemic and possibly some residual pre-leukemic cells are contained in this clone.

Patient A.6 (also shown in Figure 1e) contained mitochondrial variants which acted as main clonal markers and subclonal markers. All cells carrying the nuclear RAD21 mutation, which was observed in exome data with an allele frequency of >0.5, were assigned to this clone, again indicating that all leukemic cells are contained in the clone. These examples illustrate that CloneTracer can leverage mtSNVs called from scRNAseq for the identification of healthy/pre-leukemic and leukemic cells.

Patient A.10 constitutes a borderline case. In this patient, no nuclear mutation was efficiently captured; early drivers of the cancer were mutations in TET2 and GATA2, according to exome data. Since these mutations were not covered, we cannot be sure in what order the mitochondrial and nuclear mutations occurred, and if some of the healthy cells are instead cancerous, however, low false positive and false negative rates as determined from the presence of putatively healthy and leukemic cells in myeloid and lymphoid cells, respectively (see figure 3a,b), prompted us to include this patient into further analyses.

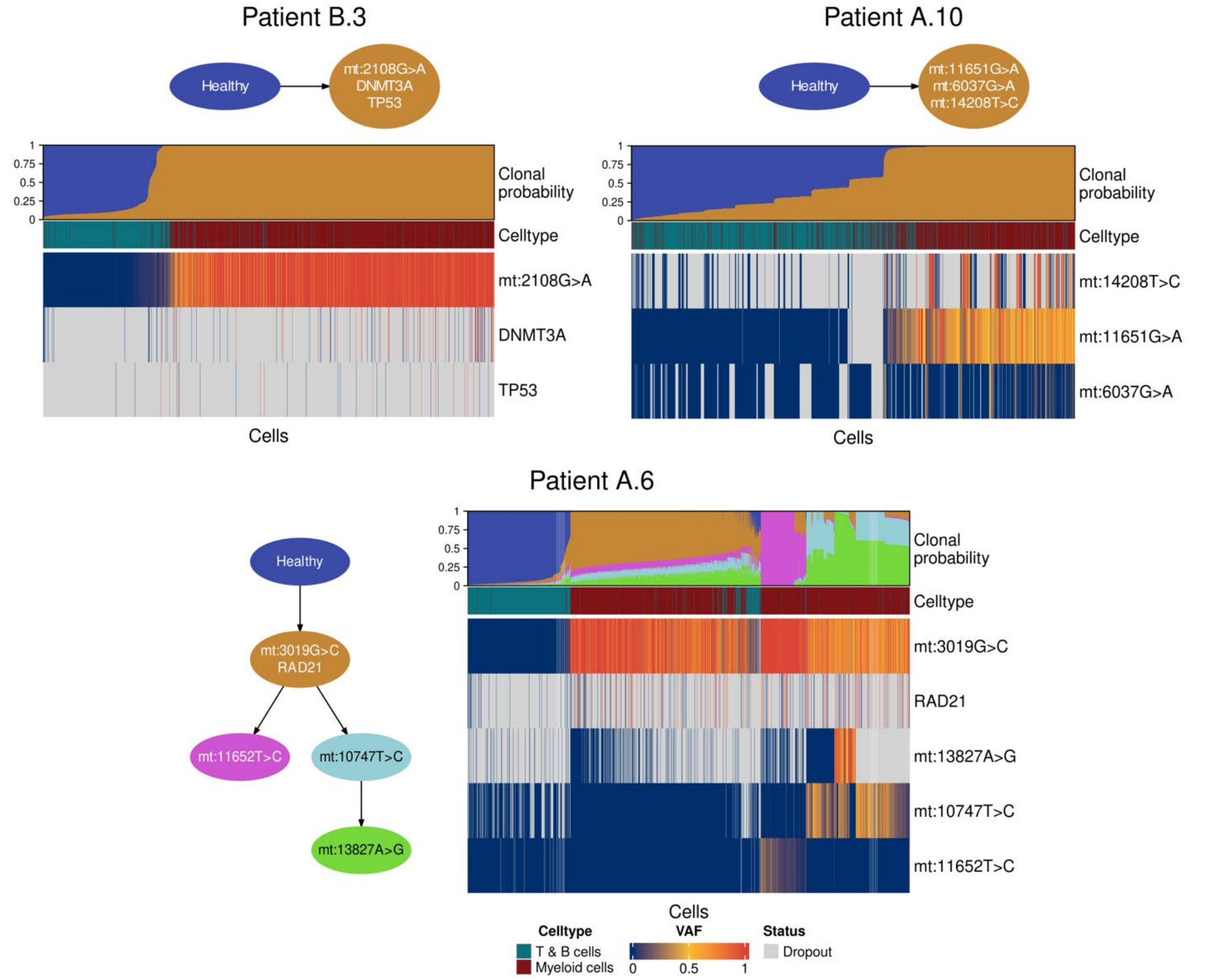

**Figure N4.** Application of CloneTracer to AML samples with mitochondrial SNVs as clonal markers. For each patient the clonal hierarchy inferred by the model is shown. The heatmaps display for each single cell the fraction of mutant reads (VAF, variant allele frequency) in all mutations as well as the cell type identity. Gray indicates that neither reference nor mutant reads were observed, termed as dropout. The clonal probabilities inferred by CloneTracer are shown at the top of the heatmap. Columns are ordered by decreasing clonal probability from the top to the bottom of the inferred clonal hierarchy.

*Samples with CNVs*

Similar to previous approaches (Gao et al., 2021, 2022), CloneTracer infers CNVs leveraging the fact that aneuploidies have an effect on the expression of genes in the affected region compared to diploid cells. In the case of large chromosomal aberrations such as monosomies or trisomies the amount of data is much larger compared to SNVs, therefore facilitating clonal tracking.

CloneTracer, unlike the aforementioned methods, requires knowledge on the location of the chromosomal aberrations. We identified large aneuploidies from clinical karyotyping of the samples. In the case of partial deletions (labelled as -part in Figure N5) we used Numbat (Gao et al., 2022) with default settings to identify the boundaries of the alterations.

The advantage that CloneTracer provides over current tools to infer CNVs from scRNAseq, is that the information on chromosomal alterations can be combined with mtSNVs and nuclear SNVs, enabling us to clarify if the CNV is really a clonal marker, and the discovery of potential subclones. This is particularly relevant in samples with a complex clonal hierarchy such as patient B.1 (Figure N5). In this sample, chromosomal aberrations in chromosomes 3 and 8 occurred early in the evolution of the leukemia and therefore are optimal markers for the distinction between healthy and leukemic cells. However, the inclusion of nuclear SNV data enabled the identification of 2 additional mutually exclusive subclones driven by mutations in *KRAS* and *NRAS* (Figure N5 and S6a).

We found other examples in which copy number alterations were part of the founding clone of the leukemia such as patients A.1 and A.3 (Figure N5). On the other hand, we also observed cases in which CNVs are subclonal as shown for patients B.4 and A.2 (Figure N5) and B.2 (Figure N6). The identified subclones are phenotypically distinct (see Figure S6d-g).

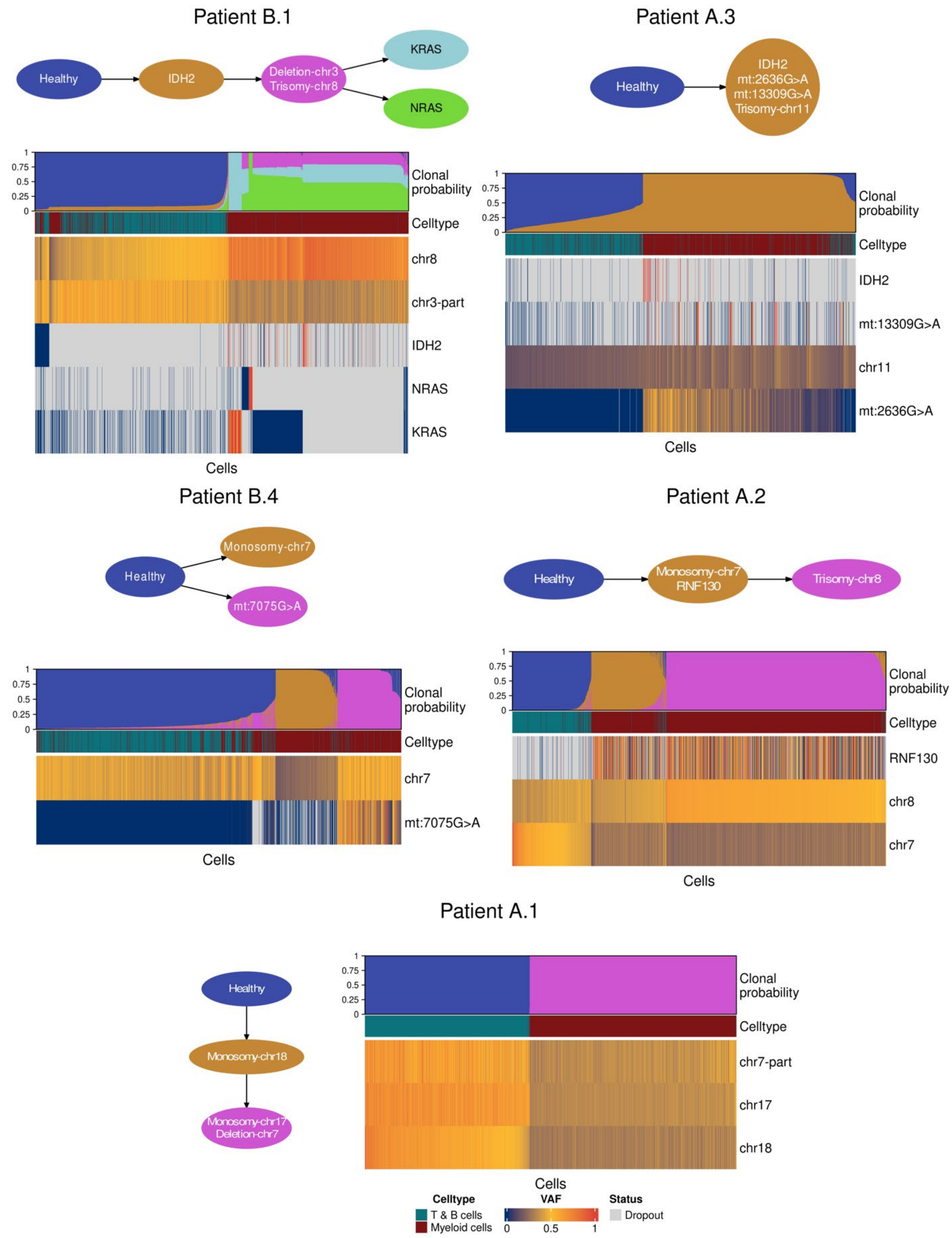

Patient B.1
Healthy
IDH2
Deletion-chr3
Trisomy-chr8
KRAS
NRAS
Clonal probability
Celltype
chr8
chr3-part
IDH2
NRAS
KRAS
Cells
Patient A.3
Healthy
IDH2
mt:2636G>A
mt:13309G>A
Trisomy-chr11
Clonal probability
Celltype
IDH2
mt:13309G>A
chr11
mt:2636G>A
Cells
Patient B.4
Healthy
Monosomy-chr7
mt:7075G>A
Clonal probability
Celltype
chr7
mt:7075G>A
Cells
Patient A.2
Healthy
Monosomy-chr7
RNF130
Trisomy-chr8
Clonal probability
Celltype
RNF130
chr8
chr7
Cells
Patient A.1
Healthy
Monosomy-chr18
Monosomy-chr17
Deletion-chr7
Clonal probability
Celltype
chr7-part
chr17
chr18
Cells
Celltype
T & B cells
Myeloid cells
VAF
0
0.5
1
Status
Dropout

**Figure N5.** Application of CloneTracer to AML samples with CNVs as clonal markers. For each patient the clonal hierarchy inferred by the model is shown. The heatmaps display for each single cell the fraction of mutant reads (VAF, variant allele frequency) in all mutations as well as the cell type identity. For chromosomal regions, the scaled fraction of reads falling into the region is shown. Gray indicates that neither reference nor mutant reads were observed, termed as dropout. The clonal probabilities inferred by CloneTracer are shown at the top of the heatmap. Columns are ordered by decreasing clonal probability from the top to the bottom of the inferred clonal hierarchy.

##### *Samples with subclonal markers*

We have shown that CloneTracer can use mtSNVs and CNVs as clonal markers for the identification of healthy and leukemic cells. However, there are instances in which these alterations occur downstream of the initial leukemic founding mutations and thereby cannot be used to distinguish healthy and malignant cells. This phenomenon occurred in patient B.2 (Figure 1f and N6) in which the mitochondrial variants and the trisomy in chromosome 8 occurred downstream of mutations in *DNMT3A* and *IDH2* genes which presumably are the initial drivers of the leukemia. In these situations, healthy cells cannot be confidently identified (Figure N8a). Notwithstanding, a large fraction of cells can unambiguously be labelled as leukemic and assigned to different subclones if present.

In the case of patient A.13, 2 mtSNVs and 3 nuclear SNVs were identified which covered between 7–44 % of cells. This enabled the reconstruction of the clonal hierarchy, however, the confident identification of healthy cells was not possible due to absence of a well-covered main clonal marker (Figure N8a). Nevertheless, a fraction of cells can be confidently assigned as leukemic.

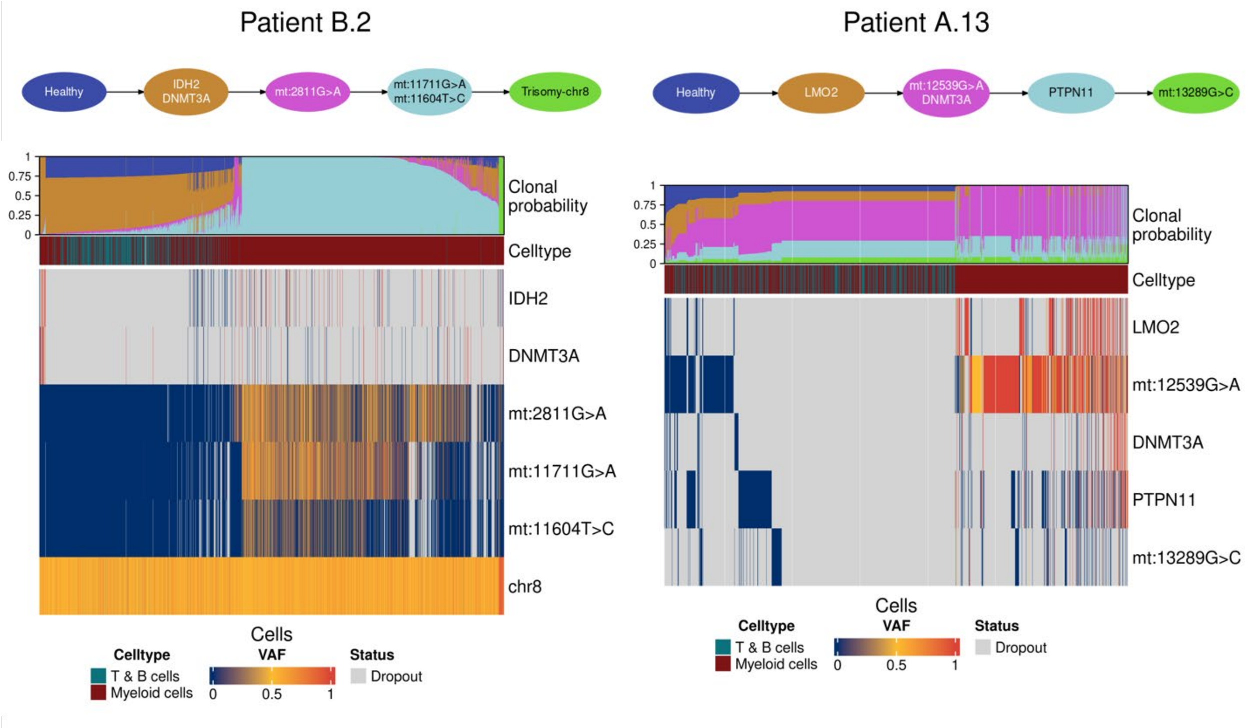

**Figure N6.** Application of CloneTracer to AML samples in which the confident identification of healthy cells is not possible. For each patient the clonal hierarchy inferred by the model is shown. The heatmaps display for each single cell the fraction of mutant reads (VAF, variant allele frequency) in all mutations as well as the cell type identity. For chromosomal regions, the scaled fraction of reads falling into the region is shown. Gray indicates that neither reference nor mutant reads were observed, termed as dropout. The clonal probabilities inferred by CloneTracer are shown at the top of the heatmap. Columns are ordered by decreasing clonal probability from the top to the bottom of the inferred clonal hierarchy.

*Samples without well-covered clonal markers*

For patients samples in which no good clonal markers were found, the inference of the hierarchy and subsequent identification of healthy and leukemic cells was not possible (Figure N7). We observed that samples in which less than 60% of cells were covered in at least one mutation CloneTracer failed (Figure N8b).

Among samples for which clonal tracking was not possible we observed cases in which neither mtSNVs nor CNVs were detected and only low-covered SNVs were amplified (patients A.4 and A.14). In other cases, mtSNVs were present but their coverage was either insufficient (patient A.7) or they clearly labelled a small subclone (patient A.15, <2% of covered cells had a mutant read and tumor bulk ATAC VAF: 0.015 for *mt:14386T>C*).

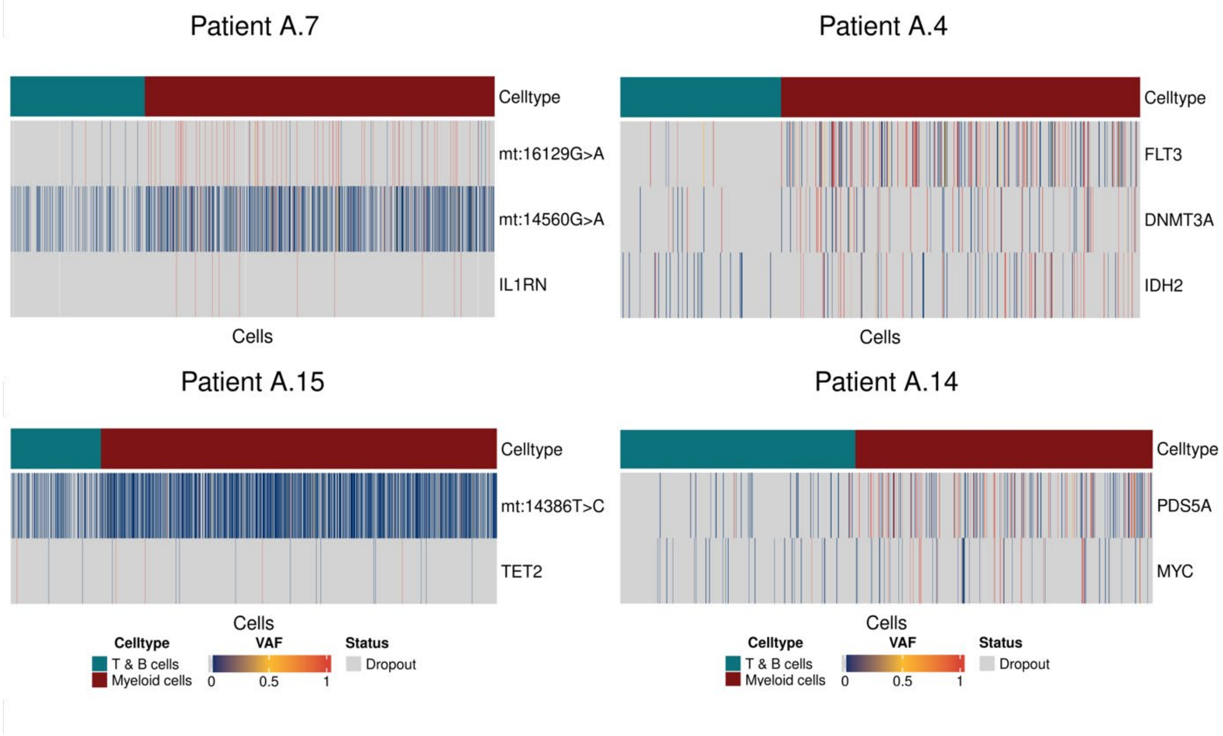

**Figure N7.** Samples without well-covered clonal markers are not suited for clonal tracking with CloneTracer. For each patient a heatmap displays for each single cell the fraction of mutant reads (VAF, variant allele frequency) in all mutations as well as the cell type identity. Gray indicates that neither reference nor mutant reads were observed, termed as dropout. Columns were order by cell type identity.

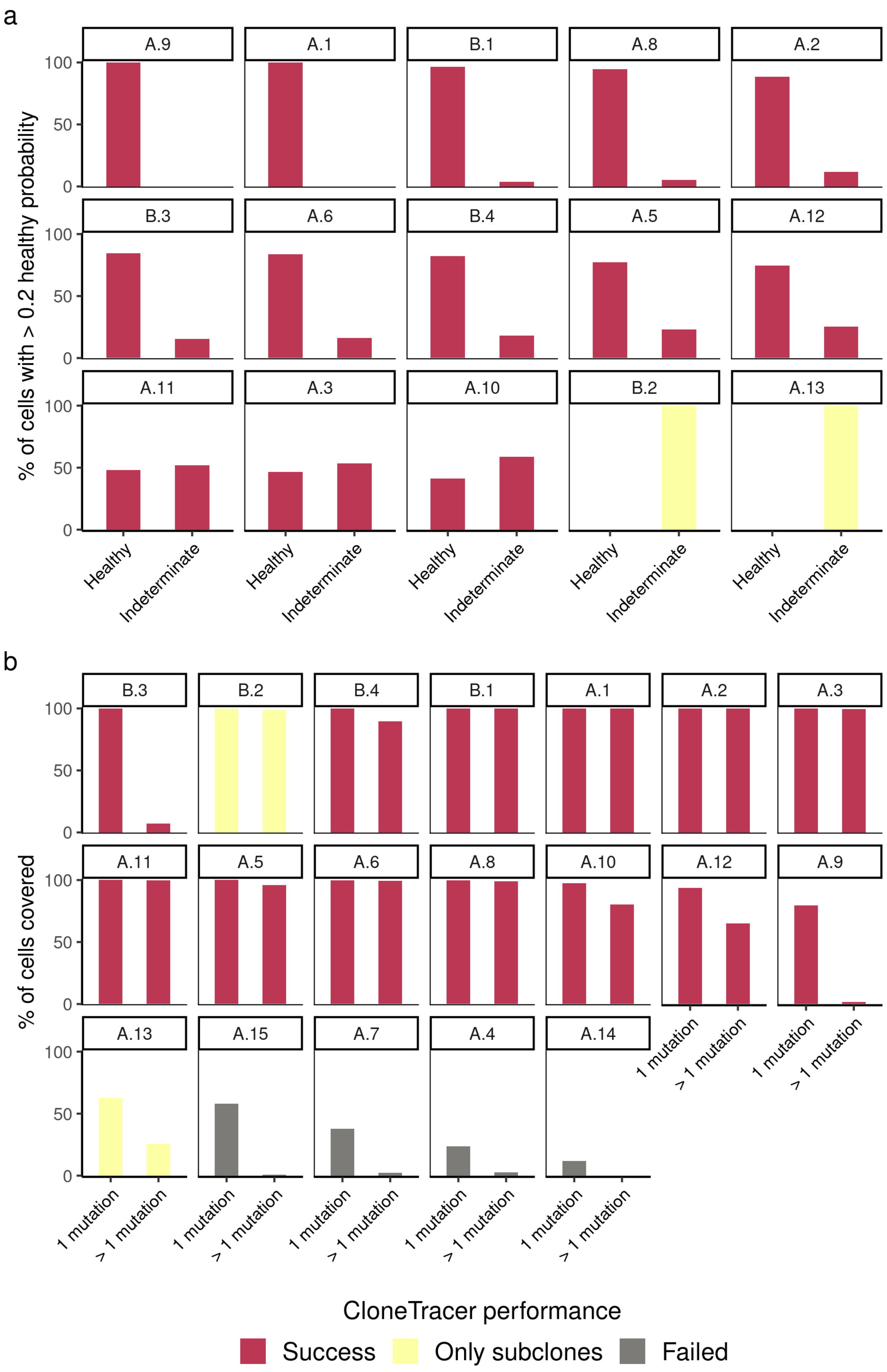
a
% of cells with > 0.2 healthy probability
A.9
A.1
B.1
A.8
A.2
B.3
A.6
B.4
A.5
A.12
A.11
A.3
A.10
B.2
A.13
100
50
0
Healthy
Indeterminate
b
% of cells covered
B.3
B.2
B.4
B.1
A.1
A.2
A.3
A.11
A.5
A.6
A.8
A.10
A.12
A.9
A.13
A.15
A.7
A.4
A.14
1 mutation
> 1 mutation
CloneTracer performance
Success
Only subclones
Failed

**Figure N8.** Criteria to determine the performance of CloneTracer. **a.** The percentage of cells assigned as healthy and indeterminate among cells with >0.2 posterior probability of being healthy is shown for each patient in which CloneTracer was ran. Samples in which the fraction of indeterminate cells was >60 % were labelled as “Only subclones”, while the rest were labelled as "Success” since healthy and leukemic cells could be confidently assigned. **b.** The percentage of cells covered in 1 mutation and at least 2 mutations is shown for each patient. Samples with <60% of cells covered in at least 1 mutation are labelled as “Failed”.
